# Origins of allostery in vertebrate hemoglobin evolution

**DOI:** 10.64898/2026.05.25.727495

**Authors:** Carlos R. Cortez-Romero, Naim M. Bautista, Collin R. Nisler, Ricardo Muñiz-Trejo, Jay F. Storz, Joseph W. Thornton

## Abstract

Allostery is an essential and structurally complex form of biochemical regulation, but how allosteric proteins evolved from nonallosteric precursors is unknown. Vertebrate hemoglobin (Hb), a symmetrical tetramer, binds organic phosphates in a central cavity between subunits, which reduces its oxygen affinity. Using ancestral protein reconstruction, we show that Hb’s historical precursor, a non-allosteric dimer, was fortuitously on the evolutionary edge of allostery: just three historical substitutions can confer allostery upon it -- one that causes tetramerization, and two others that create an effector-binding site in the cavity. The ancient tetramer could also have acquired inverse allostery – effector binding that improves oxygen affinity – via a single substitution that moves the binding site deeper in the cavity. These short evolutionary paths were possible because a key prerequisite for allostery – propensity to occupy multiple conformations that change upon oxygen binding – is an intrinsic, ancient property of the globin fold. Evolution of symmetric tetramerization caused this tertiary lability to propagate into oxygen-linked quaternary changes affecting the cavity, so the only remaining requirement for allostery was effector binding. Conformational heterogeneity and multimeric symmetry are widespread, suggesting that many allosteric proteins may have evolved by simple mechanisms from precursors fortuitously poised on the edge of allostery.

## Introduction

Allostery – change in a protein’s activity by binding to an effector molecule or other reversible modifications at locations distinct from the active site – is a critical mode of biological regulation that has been called “the second secret of life” (*1–3*). How allostery evolves is poorly understood. Allostery arises if a protein meets three necessary and sufficient conditions: it must occupy multiple conformations, those conformations must be functionally distinct, and it must bind effector with affinity that also differs between the conformations (*4–9*). Each of these features involves many amino acids, so it has been assumed that evolving allostery from a non-allosteric precursor must require mutations affecting many residues (*10–12*). Some nonallosteric proteins, however, already possess some of the prerequisites for allostery (*13–15*), raising the possibility that allostery might evolve via a few mutations that confer the remaining prerequisites (*16*).

A major challenge in understanding the historical origins of allostery is that the causes of ancient evolutionary changes in protein properties are often obscured or elaborated by subsequent modifications of sequence and structure (*17*). This challenge can be addressed experimentally by introducing historical sequence substitutions into phylogenetically reconstructed ancestral proteins expressed in the laboratory (*17, 18*). A few studies have used this strategy to understand how already-allosteric proteins evolved specificity for different effectors, and how the functional impact of effector binding in allosteric proteins may change between positive and negative regulation (*19, 20*). No work to date, however, has revealed how any protein acquired effector-driven allostery from a nonallosteric precursor during its evolution (*21*).

We used ancestral protein reconstruction to understand the evolutionary origin of allostery in vertebrate hemoglobin (Hb), the major oxygen carrier in the blood of jawed vertebrates. Hb is regulated by organic phosphates, such as inositol hexaphosphate (IHP), 2,3-bisphosphogycerate (BPG), and adenosine triphosphate (ATP), which reduce Hb’s oxygen affinity when bound and thereby facilitate oxygen delivery to metabolically active tissues in the periphery. The structural basis for allostery in extant Hb is well understood. Hb is a symmetrical α₂β₂ heterotetramer that comprises two heterodimers bound to each other isologously – using the same surface patch on each subunit but rotated 180° relative to each other. The tetramer binds four molecules of oxygen, each to a heme cofactor coordinated by one subunit. A single effector molecule is bound in a central cavity (CC), which is formed between the two β subunits (Fig. 1A) (*22, 23*). Hb can occupy two conformations, called T and R. The T conformation has lower oxygen affinity, and its CC is open; in the higher-affinity R conformation, the two α/β heterodimers are rotated relative to the other by about 15° compared to the T conformation, closing the CC (Fig. 1B). Effector binding is therefore associated with the T conformation and reduced oxygen affinity (*8, 24*).

**Fig. 1.**
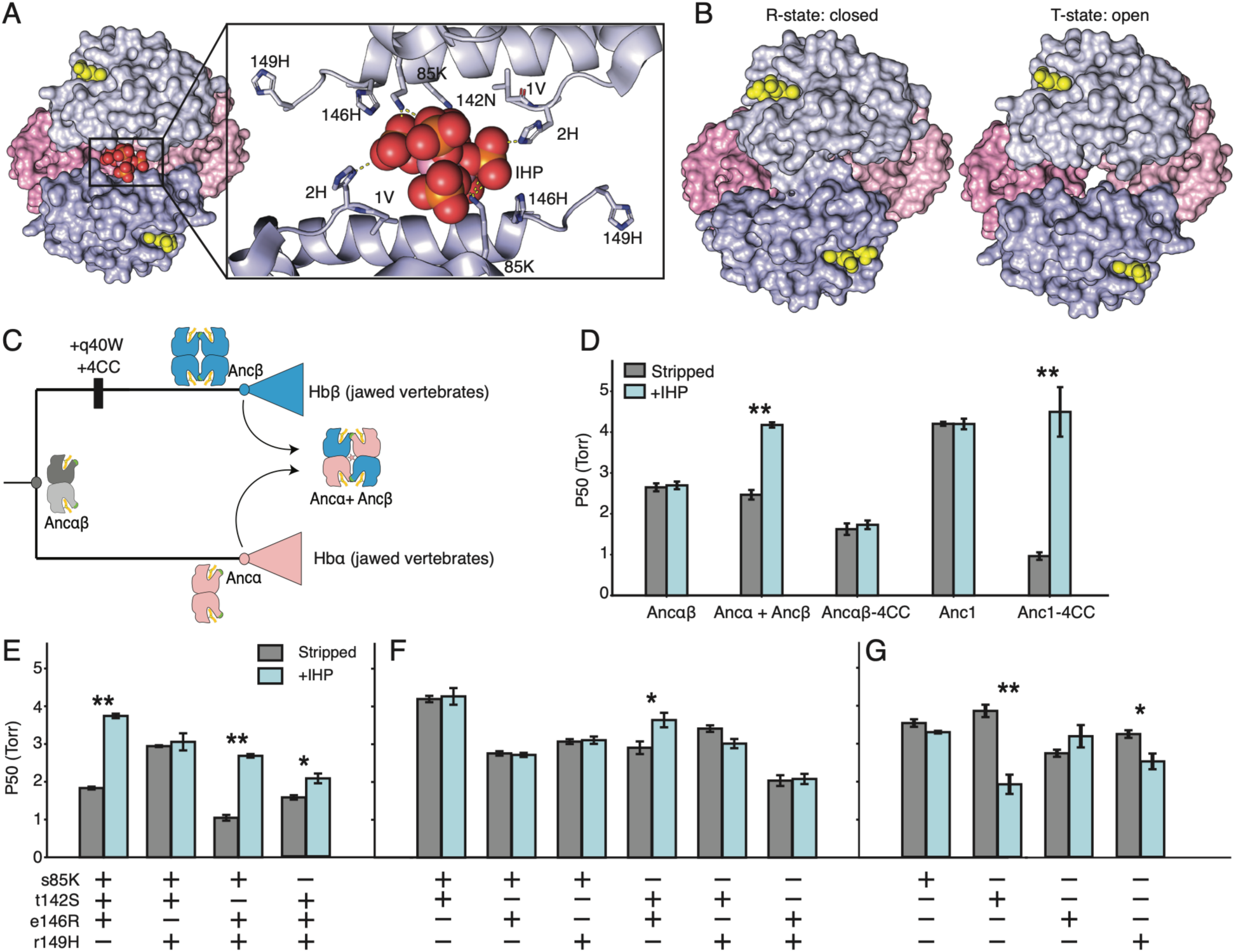
Four substitutions confer allostery in the ancestral homotetramer. **(A)** Crystal structure of Human Hb bound to IHP (PBD:3HXN). Blue, β subunits; pink, ɑ subunits; the two heterodimers that constitute the tetramer are shown in darker or lighter shades. Yellow spheres, heme. IHP is shown as spheres colored by element (red, oxygen; orange; phosphate; gray, carbons. Inset, the central cavity with IHP bound, with residue that interact with IHP shown as sticks. **(B)** *Left:* Hemoglobin tetramer central cavity is open when the hemes are deoxygenated in the absence of effector (4HHB). *Right:* The cavity is closed when hemes are bound to oxygen (2DN1). **(C)** Evolution of allosteric regulation on the HB phylogeny. Ancestral proteins and historical substitutions studied here are labeled. Icons, oligomeric states determined by prior experimental characterization of reconstructed ancestral proteins. **(D)** Allosteric effect of IHP on oxygen affinity of ancestral proteins and variants, shown as the partial pressure of oxygen at which half-maximal binding is achieved. Gray, stripped condition (IHP absent); blue, 500 μM of IHP added to stripped solution. Error bars, standard error of measurement, n = 5. **, significant difference between stripped/IHP conditions by Welch’s t-test, Benjamini-Hochberg FDR <0.05; *, FDR <0.20. **(E-G)** Oxygen affinity of all triple (E), double (F), and single (G) central cavity mutants of Anc1 in the absence and presence of IHP at 500 μM, n=3.

The phylogenetic interval during which Hb acquired allostery was previously established, but the mechanisms are unknown (*25*). The most closely related proteins to Hbα and Hbβ in the globin family are all nonallosteric monomers, including myoglobin, the major oxygen binding molecule in muscles. Our previous work showed that Hb evolved from a monomeric common ancestor with myoglobin via a homodimer (Ancαβ), which was also non-allosteric and existed just before the duplication that produced the paralogous Hbα and Hbβ lineages in jawed vertebrates. Whereas Ancαβ is non-allosteric, Ancα and Ancβ – the last common ancestral proteins of all gnathostome Hbα’s and all gnathostome Hbβ’s, respectively – assemble into a Hb-like heterotetramer that responds allosterically to organic phosphates ((*25*), Fig. 1, B and C, and figs. S1 and S2). Here we experimentally identify the genetic and biophysical mechanisms by which allosteric regulation by organic phosphates was acquired during the historical interval after duplication of Ancαβ.

## RESULTS

### Four central cavity substitutions conferred allostery

We first sought to experimentally identify the historical amino acid substitutions that caused the nonallosteric ancestor Ancαβ to acquire allostery. Because the effector binds to extant Hb tetramers in a cavity between the two Hbβ subunits, we reasoned that tetramerization would be necessary and that causal substitutions for the acquisition of allostery may have occurred at sites within the cavity during the interval between Ancαβ and Ancβ. We therefore used as a starting point Ancαβ +q40W (Anc1), which contains a single historical substitution from this same branch that was previously shown to confer high-affinity tetramerization when introduced into the ancestral dimer Ancαβ (*26*). Four amino acid changes met our criteria – three at central cavity sites that interact electrostatically with IHP’s phosphate groups (s85K, t142S, and e146R using lower and upper cases to denote ancestral and derived states, respectively) in the crystal structure of human Hb, and a fourth at C-terminal residue r149H, which occupies the cavity below IHP and makes electrostatic interactions with other residues that stabilize the T conformation (Fig. 1A). The derived states 85K and 149H are conserved among virtually all descendant Hbβ proteins, and 146R is conserved as a basic residue, whereas t142S is more variable (fig. S3).

We experimentally assessed allostery by measuring equilibrium oxygen affinity *in vitro* after stripping the medium of ionic effectors and again after adding a saturating concentration of IHP. Although the physiological effector in ancestral vertebrates is unknown, IHP serves as a useful and widely used experimental effector to test for Hb allostery (*27–30*) because it is a strong effector of Hbs from representatives of all major vertebrate lineages, whereas other organic phosphates, such as BPG, ATP, and GTP, have weaker, lineage-specific effects (*31–33*). All these effectors regulate oxygen affinity by a similar mechanism involving reversible binding to positively charged residues in the CC via their anionic phosphate groups (*23, 34, 35*).

As expected, we found that Ancαβ is non-allosteric, with an oxygen affinity of 2.6 torr, irrespective of whether IHP is present or absent (Fig. 1D). By contrast, Ancα+Ancβ is robustly allosteric, with an oxygen affinity of 2.4 torr in the stripped condition and 4.1 torr when IHP is added (Fig. 1D). The homotetramer Anc1 is nonallosteric: its oxygen affinity is 4.1 in both stripped and IHP-added conditions (Fig. 1D), indicating that tetramerization alone is not sufficient to confer allostery.

We introduced the derived states at all four central cavity sites together into Anc1 and found that the resulting protein (Anc1-4CC) is strongly allosteric, with a 4-fold difference in oxygen affinity between the stripped and IHP conditions, a stronger allosteric response than that observed in Ancα + Ancβ (Fig. 1D). Anc1-4CC is also allosterically regulated by ATP, although the response, as expected, is weaker (fig. S4).

As predicted, tetramerization is required for allostery: introducing 4CC into Ancαβ without q40W does not confer an allosteric response (Fig. 1D). Anc1 is a homotetramer, indicating that heterotetramerization is not required for allostery, as it is in Hb. Heteromerization evolved from the homomeric ancestor during the same phylogenetic interval, and we do not know the timing of this event relative to the acquisition of allostery. We found that the 4CC substitutions also confer allostery when introduced into a reconstructed ancestral Hb complex that was previously shown to assemble specifically into heterotetramers and contains no derived states in the central cavity (*25*) (fig. S5).

The evolution of allostery from a nonallosteric precursor therefore required only a few substitutions – one to confer tetramerization and at most four others at the central cavity where the effector binds. These data also show that allostery could have initially evolved in a homotetramer, so modern Hb’s dependence on the α2β2 heterotetramer for allostery was acquired later.

### Single substitutions confer inverse allostery

To understand whether all four CC historical substitutions were necessary to confer allostery, we created all possible Anc1-3CC mutants by reverting each substitution in Anc1-4CC to its ancestral state (Fig. 1E). Three of these combinations were allosteric, although the allosteric response was somewhat weaker than that of Anc1-4CC. Only e146R was necessary for allostery; this substitution and s85K, which causes the next-largest reduction in allostery when reverted, each make favorable electrostatic contacts to the effector’s phosphates in the human Hb crystal structure (*24*). Allostery can therefore be acquired by Anc1 via several different combinations of just three historical substitutions, although all four CC changes contribute to the magnitude of the response to IHP.

We next investigated whether even fewer cavity substitutions could have conferred allostery on Anc1. One double-mutant – Anc1-s142T/e146R – was weakly but significantly allosteric (Fig. 1F). Two single mutants—Anc1-t142S and Anc1-r149H—were also allosteric; however, their response to IHP was inverted, with oxygen affinity that improves when the effector is added (Fig. 1G). This inverted form of allostery is not observed in any extant organisms and is not a necessary intermediate on mutational paths from Anc1 to the derived central cavity genotype.

Taken together, these data indicate that the nonallosteric ancestral homotetramer Anc1 existed on the evolutionary edge of allostery. By just one to three substitutions that occurred during history, Anc1 can acquire either the classic mode of Hb allostery or the inverted form. Tetramerization, which is also necessary, can be conferred on Ancαβ by q40W alone, so the total number of substitutions required for the ancestral dimer to evolve into an allosteric tetramer is between two (for inverted allostery) and three or four (for the classic response).

### Ancient conformational heterogeneity predated allostery

How can so few mutations confer allostery on Anc1? Two explanations are possible. One is that the central cavity substitutions conferred all the required properties for Hb allostery -- occupancy of multiple conformations, association of these conformations with differences in oxygen affinity, and effector binding that favors one conformation over another. The other possibility is that some of these properties were already present in the ancestral tetramer for reasons unrelated to allostery. The structural location of the cavity substitutions – at the site where IHP binds, but far from the active site where heme and oxygen are bound – favors the latter hypothesis: that functionally-coupled conformational heterogeneity was already present in Anc1 before allostery evolved, and the cavity substitutions conferred effector binding at a site preferentially associated with one conformation.

We tested this hypothesis using a classic intrinsic fluorescence assay of Hb’s quaternary conformation. When extant Hb binds oxygen, the rotation of the α1β1 dimer relative to the α2β2 dimer reduces the solvent exposure of residue Trp40 at the interface between the dimers, which can be read as a reduction in fluorescence intensity at 280 nm (*36–38*). As expected, the allosteric tetramers human Hb and Ancα + Ancβ both display reduced fluorescence when oxygenated relative to when the protein solution is saturated with atmospheric nitrogen to displace oxygen (Fig. 2A; fig. S6). In the negative control experiment with dimer Ancαβ – which does not tetramerize and lacks the interfacial Trp40 residue – there is no change in fluorescence, confirming that the fluorescence signature observed in the allosteric proteins indicates quaternary change at the tetramerization interface (Fig. 2B).

**Fig. 2.**
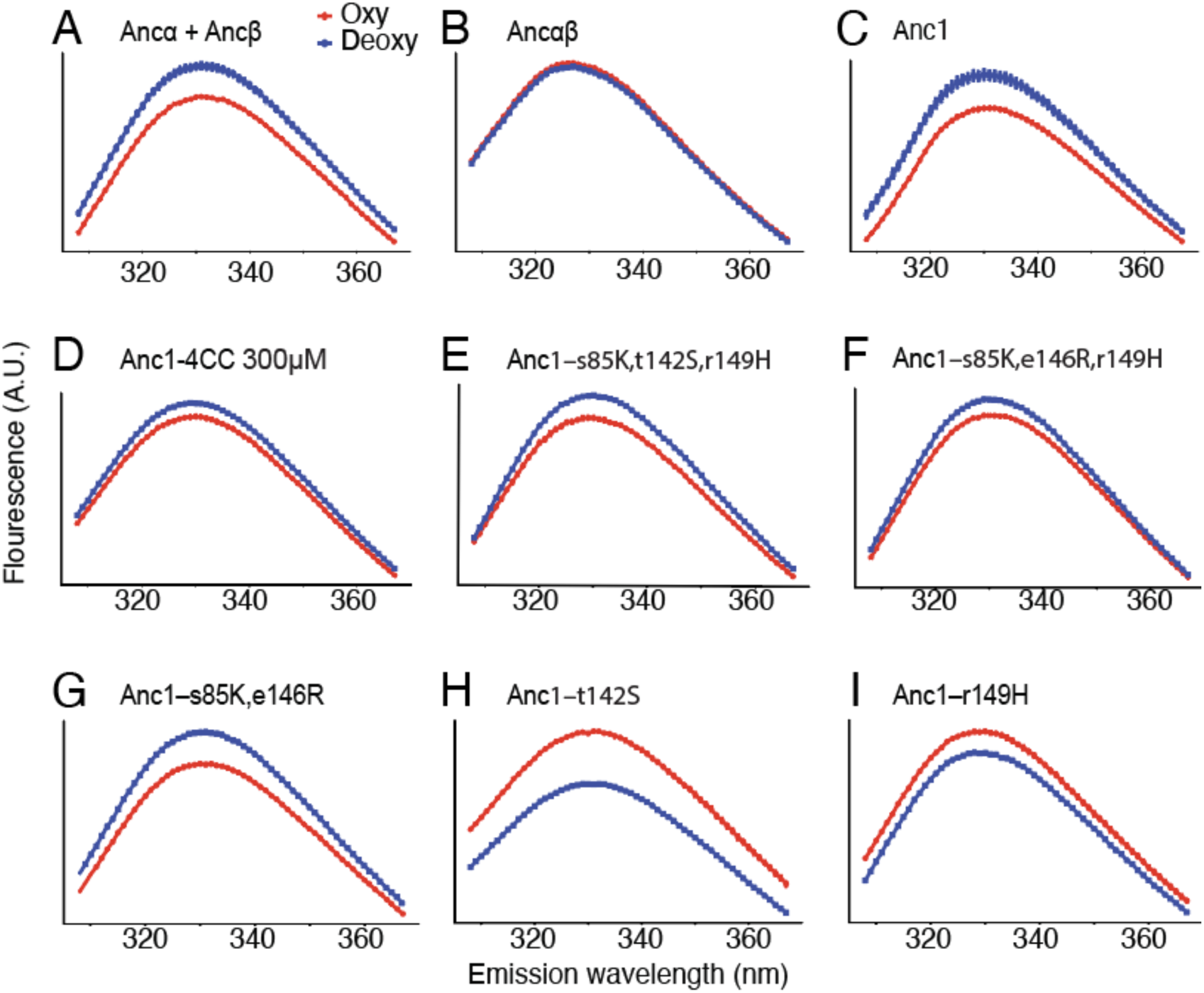
Quaternary conformational heterogeneity is present in Anc1 without allostery. **(A-I)** Fluorescence emissions scans of ancestral Hb proteins at 150 uM when excited at 280 nm. Red, oxygenated experimental condition. Blue, deoxygenated. Error bars, standard error of measurement, n = 10.

As predicted, the nonallosteric Anc1 exhibits the same reduction in fluorescence signal upon oxygenation that is observed in the allosteric Hbs (Fig. 2, A and C, and fig. S6). A similar oxygen-dependent conformational change occurred in all central cavity mutants that display the classic form of allostery, including Anc1-4CC and the allosteric triple mutants (Fig 2, D and E, and figs. S7-9). The difference in fluorescence signal upon oxygenation declines at lower protein concentrations that reduce occupancy of the tetrameric state (Fig. 2 and fig. S7). Acquisition of tetramerization alone – with no mutations in the central cavity – was therefore sufficient to confer the quaternary conformational heterogeneity required for allostery, and these conformations are differentially associated with the complex’s oxygen binding function. Thus, all requirements for allostery except for effector binding, were present in Anc1.

An inverted signal of quaternary change is apparent in the allosteric mutants of Anc1 that are inversely regulated by IHP. In these proteins, fluorescence is higher when oxygenated than deoxygenated (Fig 2H and I), indicating that Trp40 is more rather than less exposed upon oxygenation. This observation suggests that these mutations yield a novel quaternary conformation that preferentially binds to both IHP and oxygen.

### Conformational heterogeneity was mediated by ancient features

How did the acquisition of tetramerization immediately confer oxygen-mediated switching between quaternary conformations? We investigated the phylogenetic age of the major sequence and structural determinants of Hb’s tertiary and quaternary conformational heterogeneity, which are well understood in extant Hbs and involve a physical coupling between the heme pocket and the tetramerization interface (*8, 24*). Oxygen binding causes the heme cofactor to become planar, which in turn repositions two histidine residues that bind to the heme – one from “above” on helix E and one from “below” on helix F. These helices move in response, which in turn repositions secondary elements linked to those helices– a loop that connects helix E to helices D and C (the CD loop), and a turn that connects helices F and G (The FG corner). These repositioned elements are the primary contributors to the tetramerization interface, including Trp 40 (on the CD loop) and the residues on the facing subunit that interact with it (on the FG corner) (Fig. 3A). Oxygen binding thus changes the shape of the tetramerization interface, yielding different optimal orientations between facing subunits when the protein is oxygenated or deoxygenated. Because the tetramer is isologous, with the same interface repeated on all four subunits, the dimers rotate relative to each other upon oxygenation. These different quaternary orientations are stabilized by numerous other intersubunit interactions that are unique to the oxy or deoxy states (*8, 24, 39*).

**Fig. 3.**
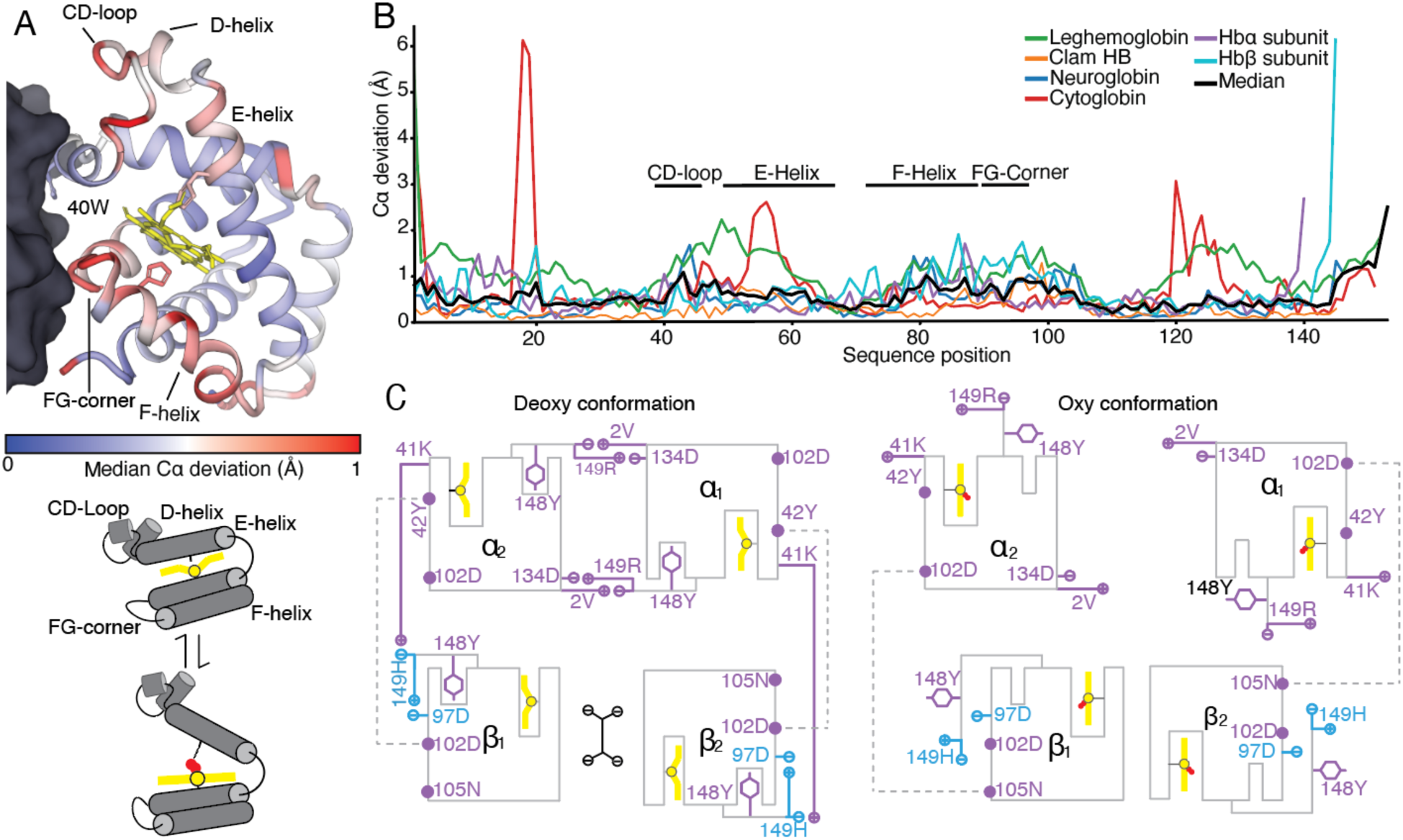
Structural causes of Hb conformational heterogeneity predated the evolution of allostery. **(A)**Tertiary change upon oxygenation at the heme pocket and tetramerization interface. *Top:* Structural relationship of heme pocket and tetramerization interface. In the structure of Anc1 predicted by AlphaFold3, one subunit across the tetramerization interface is shown as gray surface and the other as cartoon, colored by the median Cα deviation between oxygenated and deoxygenated X-ray structures of 6 globin pairs shown in panel (B). Trp40 and the histidines that bond to the heme are shown as sticks. *Bottom:* schematic of conformational change in the pocket and interface surface upon oxygen binding and release. **(B)** Deviation in the position of the Cα atom of every residue between aligned oxygenated and deoxygenated X-ray crystal structures of 6 distantly related members of the globin superfamily. Black, median across the 6 pairs. Secondary structure elements important for tertiary and quaternary heterogeneity in human Hb are marked. PDBs for deoxy and oxy structures: clam 4SDH and 1HBI, leghemoglobin 2LH7and 2GDM, cytoglobin 1V5H and 3AG0, neuroglobin 1Q1F and 1W92, human Hbα and Hbβ 4HHB and 2DN1. **(C)** Reproduction of Perutz’s classic diagram of electrostatic interactions distinguishing the deoxygenated (left) and oxygenated (right) quaternary conformations. Single-letter amino acid code shows the states present in Ancα+Ancβ, colored by their phylogenetic age: purple, states that evolved prior to Ancαβ; blue, residues substituted on the branch leading to Ancβ. Yellow, heme; red, oxygen; IHP, black. Residue 2V is the N-terminal residue of Hb following cleavage of the initial methionine.

Structural analysis in a phylogenetic context establishes that these features evolved long before Ancαβ and the acquisition of allostery or tetramerization. We assessed the history of tertiary heterogeneity by aligning pairs of oxygenated and deoxygenated X-ray crystal structures of several distantly related globin family members that are nonallosteric monomers, including globins from mollusks and plants; for each protein, we then measured the difference in Cα position of each residue between the oxygenated and deoxygenated structures. In every case, we found major conformational differences at the CD-loop, E-helix, F-helix, and FG-corner (Fig. 3B, and fig. S10); this result extends previous work that had documented ligation-induced movements in the same regions of several homologous globins, including myoglobin, neuroglobin, cytoglobin, and the blood globins of agnathans (*40–45*). These oxygen-mediated tertiary motions have therefore been present since at least the last common ancestor of animals and plants, ∼1 billion years prior to the evolution of Hb allosteric regulation in jawed vertebrates. This ancient heterogeneity can be explained by the fundamental chemistry of heme – which intrinsically changes planarity when oxygenated – and by the tertiary fold that is common to all globins: in all these proteins, the heme is coordinated by homologous histidine residues on homologous helices E and F, which lead to the same patch on the surface (Figs 3B, S10), but only in vertebrate Hb is this patch used for tetramerization.

At the quaternary level, the residues that mediate the distinct contacts that stabilize the oxygenated and deoxygenated conformations in modern Hb also predated the evolution of allostery. Figure 3C reproduces Perutz’s classic diagram of electrostatic interactions that distinguish Hb’s T and R conformations but adding information about the age of the residues. Of 7 interactions that are distinct between the human Hb T and R conformations, 6 involve residues that were all acquired before Ancαβ.

The conformational prerequisites for Hb allostery and its sequence and structural determinants are therefore ancient, substantially predating the evolution of allostery. As soon as tetramerization was acquired, ancient tertiary heterogeneity propagated into distinct oxygen-dependent quaternary rotations, setting the stage for the rapid acquisition of allostery when effector binding evolved at the central cavity.

### Structural mechanisms for the evolution of allostery

Finally, we used molecular dynamics simulations to gain insight into potential structural mechanisms by which the historical cavity substitutions caused allostery to evolve. *Creating an effector binding site.* The simplest hypothesis to explain how the CC substitutions confer allostery on Anc1 is that they introduce residues that directly facilitate binding of the effector at a site that is accessible only when the protein is in its deoxygenated state. To test this hypothesis, we initiated MD trajectories from predicted structural models of Anc1 and Anc1- 4CC tetramers containing deoxygenated heme and IHP bound in the central cavity at the same location as in human Hb (fig. S11). Across a 400 ns trajectory, IHP remained stably bound at this site in Anc1-4CC (Fig. 4, A and B, and fig. S12A), but it was ejected from Anc1’s cavity within ∼20 ns (Fig. 4, C and D, and fig. S12B).

**Fig. 4.**
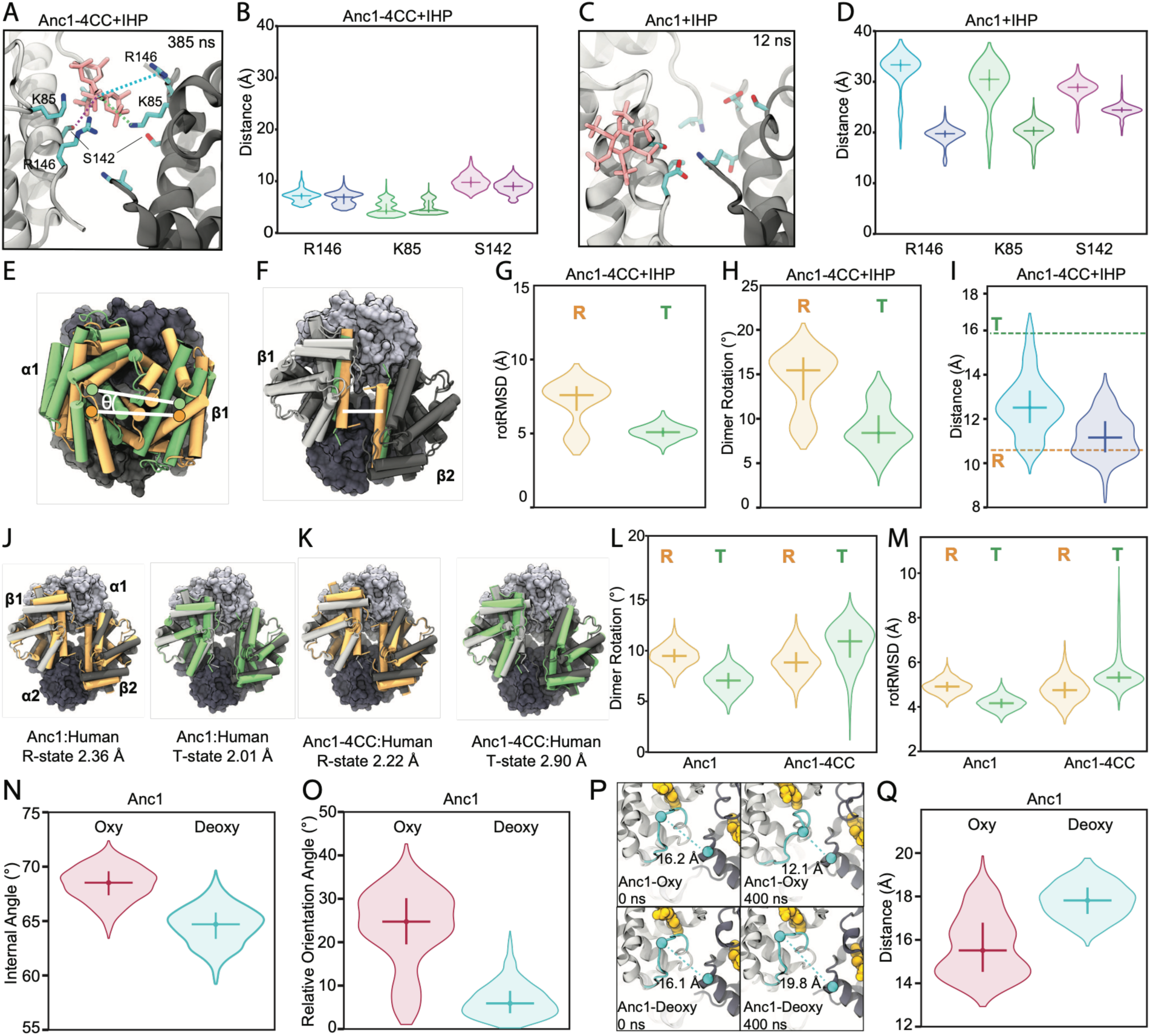
Structural mechanisms for the evolution of allostery. **(A-D)** Interactions between IHP and central cavity residues in Anc1-4CC **(A,B)** or Anc1 **(C,D)** in molecular dynamics simulations. **A)** Structure at the indicate timepoint, showing stable coordination of IHP. Dotted lines, distances measured and plotted across each trajectory, colored as in panel B. Each residue appears twice in the pocket and its distance was measured separately. Pink sticks, IHP; cyan, 4CC substituted residues. Backbones of the two subunits are in different shades of gray. Distances were measured from the center-of-mass of IHP to the terminal O and N atoms of Ser and Lys and from Cζ of Arg. **B)** Distribution of distances to IHP across the trajectory (initial 20 ns removed). Horizontal line, median; vertical line, interquartile range. **C,D)** Same as panels A and B but for Anc1, showing loss of contact with IHP early in the trajectory. **(E)** Metrics of difference in quaternary structure between two Hb tetramers, using the human Hb R (orange) and T (green) states as an example (PDBs 1RD and 2HHB). Two tetramers are aligned to each other by minimizing the RMSD between Cα atoms in one of the dimers from each tetramer (gray surfaces); the unaligned dimers are shown in orange and green. Circles, center of mass of the subunits within each unaligned dimer, with the vector between them shown as a white line. θ, dimer rotation angle between these vectors. rotRMSD is the RMSD between Cα atoms of the unaligned dimers. Subunits are labeled. For both metrics, lower values indicate more similar quaternary structures. **(F)** Distance between the H- helices across the central cavity (white line segment). **(G,H)** Distribution of rotRMSD and rotation angle for Anc1-4CC-IHP when aligned to either the human Hb R (orange) or T state (green). **I)** Distribution of interhelical distance for Anc1-4CC-IHP. Distances between the two pairs of H-helices in the homotetramer are plotted separately. Dotted lines, distance in human R-state and T-state structures. **(J)** Equilibrated AlphaFold3 model of Anc1 without IHP (gray) is shown aligned to the crystal structures of the human R-state (left, orange) or T-state (right, green). Alignment and RMSD are for Cα atoms. **(K)** Same as panel J, but for Anc1-4CC without IHP. **(L.M)** Distribution of dimer rotation angle (L) and rotRMSD (M) for Anc1 and Anc1-4CC in the absence of IHP relative to human R and T states. **N-O)** Differences in quaternary structure of Anc1 with oxygenated vs. deoxygenated heme. The plots show the distributions of internal angle and relative orientation angle (metrics of relative rotation between two dimers within a tetramer, as defined in fig S19). **P,Q)** Differences in the tetramerization interface between oxy- and deoxy-Anc1. **P)** Structural snapshots of the interface and heme pocket are shown at designated timepoints. Dotted line, distance across the interface from the CD loop of one subunit to the FG corner of the other subunit (measured between Cαs of H47 and K98, dots). Q) Distribution of distances from CD loop to FG corner across the trajectories, as shown in panel P.

As predicted, the four cavity substitutions make the pocket electrostatically and sterically complementary with IHP. The derived residues Lys85 and Arg146 are positively charged, and they form stable electrostatic interactions with IHP’s negatively charged phosphate groups (Fig. 4, A and B); by contrast, the ancestral residues at these sites in Anc1 (glu85 and glu146) are negatively charged and repel the effector (Fig. 4, C and D, and figs. S12 and S13). Ser142 in Anc1-4CC forms transient hydrogen bonds with IHP (Fig. 4A) and removes a steric clash between the effector and the extra methyl group on the Anc1’s thr42 (Fig. 4, A and B). The fourth substitution (r149H at the protein’s C-terminus) relieves a major steric clash with the effector: the ancestral arg149 occupies the central cavity, where it is stabilized by a salt bridge to Asp134, but substituting this residue to His149 in Anc1-4CC compromises this interaction, allowing the C-terminus to shift outward and making space for IHP (fig. S14). Because the Anc1-4CC tetramer is symmetrical, each derived residue appears twice in the binding pocket – once on each subunit – so each favorable interaction occurs twice in Anc1-4CC, along both sides of IHP (Fig. 4A).

Also as predicted, IHP binding in Anc1-4CC is preferentially associated with a low-oxygen affinity quaternary conformation. We quantified the quaternary similarity of Anc1-4CC to the classic T and R states of human Hb (which have low and high oxygen affinity, respectively) by aligning Anc1-4CC to each human X-ray crystal structure using only one of the dimers in each tetramer and then measuring the angle of rotation and the Cα RMSD between the two unaligned dimers (Fig. 4E). We also measured the distance between the H-helices of the two subunits across the central cavity, which is much larger in the human T than R conformation, allowing the former to accommodate the effector (Fig. 4F). In the trajectory with IHP, Anc1-4CC undergoes a notable transition in quaternary structure, becoming much more similar to the human T conformation by these metrics, and the interhelical distance increases by about 2.5 Å, allowing sufficient space for the effector (Fig. 4G-I and fig. S15). This conformational transition is not observed in the trajectory of Anc1 (fig. S16) and is therefore a consequence of IHP binding by Anc1-4CC. The 4CC substitutions therefore conferred allostery by generating a complementary binding surface for the effector that is uniquely accessible in the deoxygenated state.

#### Increased oxygen affinity in the absence of IHP

The 4CC substitutions improve oxygen affinity in the absence of IHP, magnifying the allosteric response to the effector. A simple mechanism to explain this observation would be that the 4CC mutations increase the propensity of the protein to occupy an R-like state when IHP is absent. Corroborating this hypothesis, the quaternary conformation of Anc1-4CC without IHP favors an R-like state much more than Anc1 (Fig. 4J-M and fig S17). This change arises because the 4CC substitutions abolish interactions that favor the T-like conformation of Anc1, thus reducing occupancy of the state (Fig. 4L and, 4M).

Specifically, ancestral residues glu146 and arg149 in Anc1 form strong electrostatic interactions across the central cavity in the T conformation; ser85 also hydrogen bonds to either glu146 on the same subunit or His3 on the opposing subunit (fig S18). Substitution of these residues to the derived states abolishes these interactions and thus destabilizes the deoxy conformation in Anc1- 4CC when IHP is absent (fig S13).

#### Ancient conformational heterogeneity

To gain insight into the mechanisms underlying the ancient oxygen-dependent conformational heterogeneity of Anc1, we compared MD simulations of this protein when the heme is oxygenated or deoxygenated. Beginning from the equilibrated T-like structure of Anc1-deoxy described above, we removed the IHP, and initiated 400 ns MD simulations, using either oxygenated or deoxygenated heme. We assessed potential changes in quaternary structure during this trajectory using two metrics of quaternary structure: first, the angle of rotation between the central axes of the two dimers within each tetramer (which captures the overall orientation of the two dimers relative to each other) and second, the angle of rotation between a vector defined by the dominant principle axis of the H-helices in one dimer and that in the other (which captures the impact of quaternary rotation on the helices that frame the central cavity, Fig. 4N,O; fig S19A & B).

We observed a quaternary transition in the presence of oxygen. When the heme is deoxygenated, the central axis angle of Anc1 increases slightly, and the H-helix rotation angle decreases as the protein adjusts to the removal of IHP from the central cavity. By contrast, the trajectories with oxygenated heme show a marked reduction in the central axis angle and a ∼15 degree increase in the helix rotation angle (Fig. 4N,O). These differences in oxy-vs-deoxy conformations are qualitatively similar to those between the human Hb R and T conformations, although the magnitude of difference in the ancestral protein is smaller (fig. S20).

The quaternary changes caused by oxygen binding in Anc1 are associated with tertiary changes near the heme pocket that change the multimerization interface and favor the transition to a new quaternary conformation. In the starting deoxy structure, the loop between helices C and D is close to the heme binding site; with oxygenated heme, this loop flips away from the bottom edge of the heme (Fig. 4P,Q; fig S19C). These changes alter the network of interactions across the interface (Fig. S21). The quaternary rotation of Anc1 coupled to oxygen binding is therefore associated with tertiary changes in the heme pocket and tetramerization surface that are qualitatively similar to those observed in extant Hb.

Taken together, these data provide a structural explanation for the acquisition of allostery when a few central cavity substitutions are introduced into the nonallosteric precursor Anc1 (summarized in Fig. 6). As soon as Trp40 conferred tetramerization on Anc1, ancient tertiary heterogeneity was transformed into oxygen-linked quaternary heterogeneity, with a structural basis that is similar to that in modern Hb. All that remained was for IHP binding to be acquired at a site preferentially available in the low-oxygen-affinity conformation – precisely what that the 4CC substitutions do.

### Mechanisms for evolution of inverse allostery

Finally, we used MD simulations to understand how inverse allostery is conferred on Anc1 by single central cavity substitutions. We focused on Anc1-t142S, because this mutant had the strongest inverted response. We used Alphafold3 to generate an initial structure of this protein with oxygenated heme, equilibrated it for 100 ns, docked IHP into the structure, and initiated a 300 ns trajectory.

As predicted by the fluorimetry experiments, the Anc1-t42S mutant occupies a different conformation because it binds IHP in a novel way. Across the entire trajectory of this complex, IHP is stably bound, but at a location that equilibrates ∼5 Å deeper in the cavity than its canonical position in Anc1-4CC-IHP and human Hb (Fig. 5A & B; fig S22). IHP is stabilized in this new location by hydrogen-bonds to S142, as well as salt bridges to the ancestral residues arg149 and the N-terminal amine, which are pulled into the central cavity (Fig. 5C, fig S23, S24A & B). These latter interactions are not observed in any other ancestral proteins we examined or the human Hb crystal structures. They are specifically attributable to mutation t142S, because a control simulation of Anc1 beginning from the exact same structure – but with S142 reverted to threonine – did not retain IHP in the cavity (fig. S25).

**Fig. 5.**
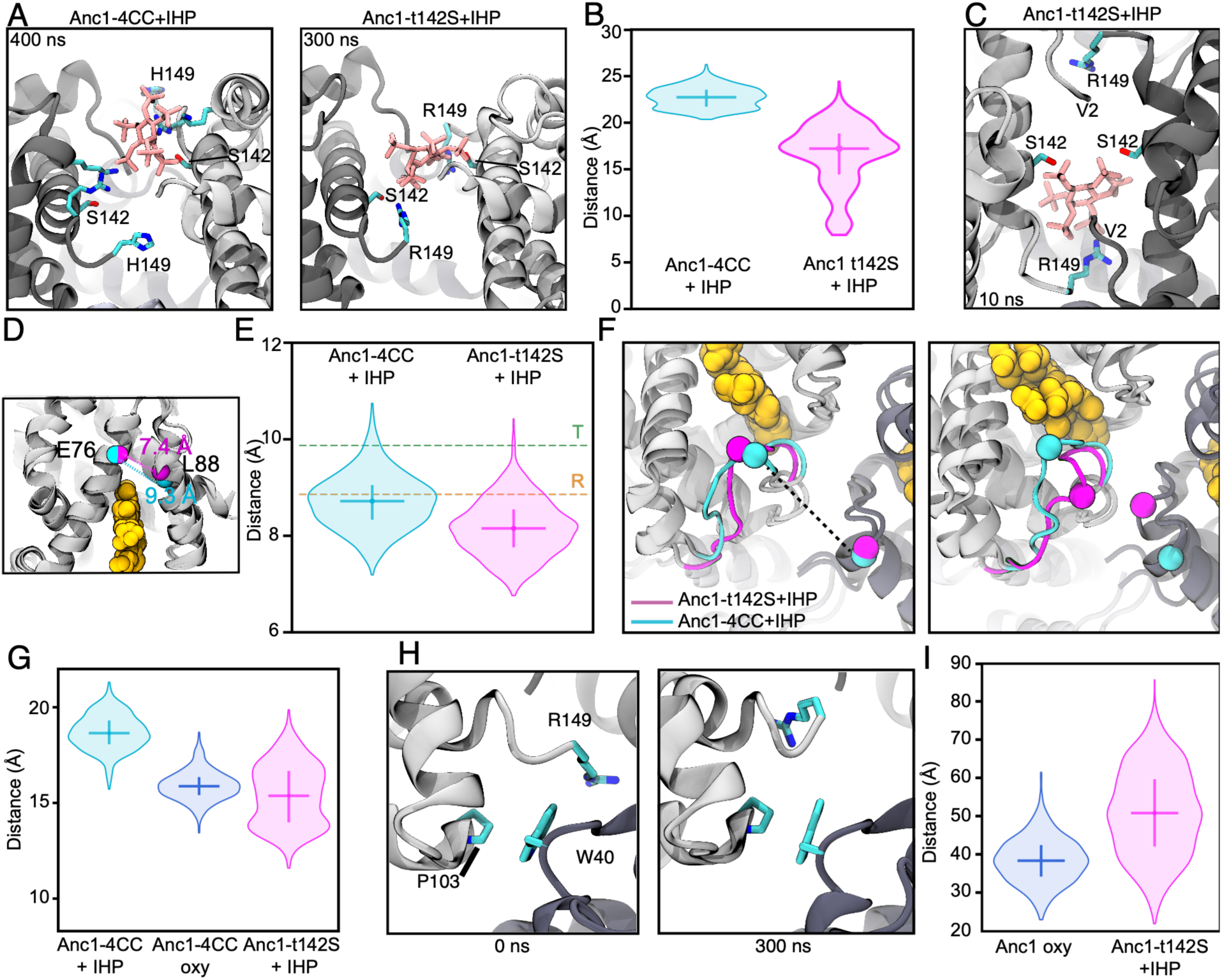
Mechanisms of inverted allostery in Anc1-t142S. **(A)** Binding of IHP in Anc1-4CC (*left*) and Anc1- t142S+oxy (*right*), showing the deeper location of IHP in the latter case and its close association with S142 and the C-terminal R149. **(B)** Distance between the center of mass of the tetrameric multimer and the center of mass of IHP across the trajectories of Anc1-4CC- IHP and Anc1-t142S+oxy. See also fig. S22. **(C)** Close view of the central cavity of the Anc1-t142S model with docked IHP after equilibration, showing N- and C-terminal residues V2 (after cleavage of initial Met) and R149. **(D-E)** Tertiary differences in the heme pocket. **D)** The distance from helix E to helix F was measured using the Cα atoms of E76 and L88 (spheres) as landmarks. Yellow spheres, heme. A superposition is shown of these helices and atoms from an exemplary frame in trajectories of Anc1-4CC-IHP-deoxy (cyan) and Anc1-t142S-IHP-oxy (pink).**E)** Distribution of the distance between helices E and F (mean across the four tetramers, measured as in panel D) for each trajectory. Dotted lines, distance for human R- and T-state crystal structures (PDB 1IRD and 2HHB). **F-H)** Tertiary differences at the tetramerization interface. **F)** Distance between the CD loop and the FG corner of its neighboring subunit, as described in Fig. 4P. Structures from representative frames of each trajectory are shown. **G)** Distribution of CD-to-FG distances (mean across four subunits) across trajectories of Anc1-t142S-IHP+oxy and Anc1-4CC. **(H)** Change in interactions at the interface of Anc1-t142S-IHP. The C-terminal carboxyl of R149 is pulled away from Trp 40, as shown in structures from the beginning (left) and end (right) of the trajectory. **I)** Increased solvent-accessible surface area of Trp40 residues in Anc1-t142S-IHP+oxy is greater than that of Anc1+oxy, consistent with its increased fluorescence when oxygenated.

The novel quaternary conformation of Anc1-t142S-oxy are associated with tertiary differences that explain the protein’s inverted allostery. Pulling the C-terminus into the central cavity to interact with IHP is associated with a change in the F helix, which packs against the C-terminus on one side and moves inward towards the heme by ∼1 Å (Fig. 5D & E; fig S24D & E, fig S26). This narrowing of the heme pocket is similar to that observed in the R-state of human hemoglobin. In addition, the C/D loop in the oxygenated Anc1-t142S-IHP complex flips away from the heme’s bottom edge, packing in a way that is more similar to Anc1-4CC’s oxy than deoxy conformation (Fig. 5F,G, and fig. S27). These observations provide a likely mechanism for the increased oxygen affinity associated with IHP binding by this protein.

Finally, these changes in Anc1-t142S affect the tetramerization interface, explaining why Anc1- 142S displays an inverted fluorimetry pattern. In Anc1 and human Hb, residues on the C-terminal helix pack around Trp40, but repositioning this helix in Anc1142S-IHP-oxy compromises these interactions and increases solvent exposure of Trp40 (Fig. 5H and I, and fig. S28). Our finding that substitution r149H – at the protein’s C-terminus – also confers inverted allostery and fluorimetry is consistent with the key role of the C-terminal helix in mediating this behavior.

Taken together, the MD simulations provide a detailed explanation for how introducing CC substitutions in various combinations into Anc1 can confer allostery in the classical or inverted form. Different combinations of derived and ancestral residues cause IHP to bind at different locations, which are associated with different tertiary adjustments. These differences affect the heme binding pocket, determining whether oxygen binding is favored or disfavored when the effector is bound.

## DISCUSSION

Our work shows that Ancαβ, the nonallosteric dimer that historically preceded the allosteric Hb tetramer, was near the evolutionary edge of evolving allosteric regulation. Allostery in either the classical or inverted forms could be conferred on this protein by several different combinations of just a few historical substitutions. These short paths to allostery were possible because key structural requirements for allostery were already present in the non-allosteric precursor Ancαβ as biochemical spandrels –by-products of protein structure that was initially functionally inconsequential but was eventually incorporated into allosteric regulation (*46*). Heterogeneity of tertiary conformation linked to oxygen binding arose directly from the basic globin fold and the fundamental chemistry of heme. When tetramerization in Hb evolved at a surface that was conformationally coupled to the heme pocket, this tertiary heterogeneity immediately propagated into a large-scale quaternary rotation that affects the central cavity’s availability. Evolving effector binding in this cavity allowed the effector to act as a wedge that favors the conformation with low oxygen affinity (Fig. 6).

**Figure 6.**
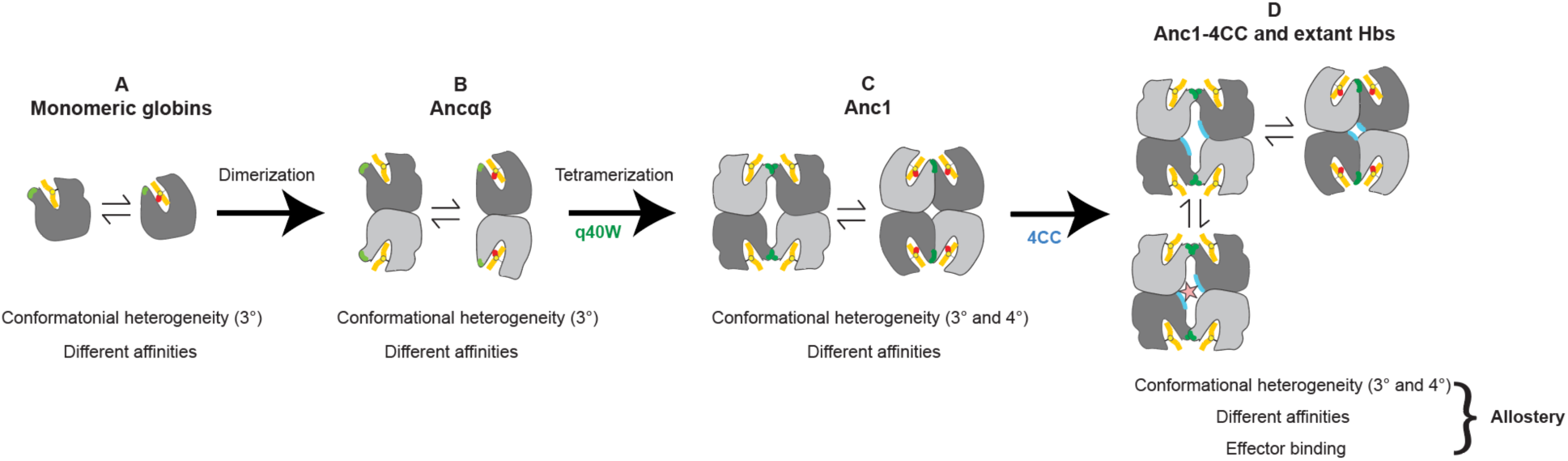
Evolution of features required for Hb allostery. **(A)** Ancient monomeric globins occupy at least two conformations that differ in tertiary structure around the pocket for heme (yellow) and at the surface that later became the tetramerization interface (light green). They also differ in affinity for oxygen (red). (**B)** These features were retained in the dimeric Hb ancestor Ancαβ. (**C)** Substitution Trp40 (dark green) at the interface surface conferred tetramerization. The tertiary conformations are oriented across the interface in differently rotations, so the tetramer occupies multiple quaternary conformations that differ in the accessibility of the central cavity and in oxygen affinity. (**D)** Adding the 4CC substitutions (blue) creates a binding site for IHP (pink star) that is preferentially accessible in the low-affinity conformation. The historical order of the Trp40W and 4CC is unresolved; one possible order is shown here.

Our results suggest that the prerequisites for allostery were acquired contingently rather than via long paths deterministically driven by selection for allosteric regulation. If tetramerization had never evolved, or if it had evolved at a different patch that is not conformationally linked to the remodeling of the heme binding site, then allostery could not have evolved by the substitutions in the central cavity. The prerequisites that poised Hb’s evolutionary precursors at the edge of allostery could not have been acquired because of selection for allostery, because they were not sufficient to confer even a weak allosteric response. It has been proposed that the conformational heterogeneity that underlies allostery may evolve because of selection on proteins to be adaptable in the face of fluctuating biological environments (*47*). In hemoglobin, at least, this does not appear to be the case: ancient tertiary heterogeneity requires no selective explanation because it has no known functional consequences and arises as a simple by-product of the protein’s architecture. Our data do not resolve the order in which the key substitutions occurred, but whether q40W or the CC substitutions occurred first, neither of these intermediate states would have been allosteric, but both would have been just one or two substitutions away from acquiring allostery. In either case, it is possible that the final substitution that conferred allostery could have increased fitness, but all those that occurred earlier could not have been fixed by selection for allostery, because they were not sufficient to confer it, even weakly.

Features similar to those that allowed Hb’s precursor to suddenly acquire allostery are widespread among proteins; it therefore seems likely that many other allosteric proteins also evolved via short paths from nonallosteric proteins that were fortuitously poised on allostery’s edge. Like globins, virtually all proteins occupy multiple conformations rather than a single rigid structure (*48–50*). These conformations typically differ in activity, as implied by induced fit and conformational selection mechanisms for ligand binding and conformational mobility as the basis for catalysis (*51, 52*). Coupling of conformational change in the active site to change at other locations is also nearly universal, simply because a protein’s fold involves covalent linkages, dense packing between elements, and regions with many degrees of freedom (*53–55*). In any protein with such features, acquiring effector binding at any coupled site can confer allostery. It has been established that binding to small molecules or proteins can often be conferred by one or a few mutations on a protein’s surface (*16, 56*). Supporting the idea that proteins are often on the evolutionary edge of allostery, biochemical studies have shown that allostery can be engineered into extant nonallosteric extant proteins by single point mutations or insertions of effector-binding sites (*14, 57–60*). Even some proteins that have no known endogenous allosteric regulators are “latently allosteric:” they can be regulated by drugs or other substances that happen to bind to surface sites conformationally associated with the active site, indicating that they possess all the prerequisites for allostery without any natural biological function (*13, 61, 62*).

Across the protein universe, allostery is strongly enriched among multimers with isologous architecture (*26, 63–65*), and our findings suggest why this may be. First, we found that isology facilitated the acquisition of effector binding by multiplying the impacts of favorable substitutions, each of which appears multiple times within the symmetrical central cavity, doubling its energetic contribution to binding and exponentially increasing its effect on affinity (*26*). Second, isologous multimerization dramatically enlarged the set of sites at which acquiring effector binding could have conferred allostery. Tertiary heterogeneity in globins is linked to a relatively small surface where tetramerization evolved, but the quaternary rotation that results from isologous multimerization affects not only the central cavity but all surfaces that comprise multiple subunits in the tetramer. If effector binding had evolved at any such site, allostery would have arisen. Isologous multimers are therefore more likely to be at the evolutionary edge and to have more mutational opportunities to acquire allostery.

Isologous multimers themselves can easily evolve by one or a few substitutions at various locations on a protein’s surface (*25, 26, 66*). Our data suggest that all such protein complexes, and many monomers too, are often at or near the edge of acquiring allostery. The particular structural and functional form of allostery that evolves – binding to this or that effector, here or there on the protein’s surface, enhancing or impairing its biological activity -- depends on the past mutations that brought it to the edge and on those that happen to push it over into the future.

## METHODS

### Sequence data, alignment phylogeny, and ancestral sequence reconstruction

The reconstructed ancestral sequences used here are the same as those reported previously (*25*), with Genbank IDs MT079112, MT079113, MT079114, and MT079115. In brief: 177 amino acid sequences of hemoglobin and related paralogs were collected and aligned. The maximum likelihood (ML) phylogenetic tree was inferred using the AIC best-fit model, LG+G+F (*67*). The phylogeny was rooted using as outgroups neuroglobin and globin X, which are found in both deuterostomes and protostomes and diverged prior to the gene duplications that produced vertebrate myoglobin and the hemoglobin subunits. Ancestral sequence reconstruction was performed using the empirical Bayes method, given the alignment, ML phylogeny, ML branch lengths, and ML model parameters (*68*). For details, see ref. (*25*). The historical central cavity mutations tested here include all four residues substituted between the maximum a posterior sequences of Ancαβ and Ancβ at sites that are within the central cavity in the crystal structures of human Hb (e.g., PDB IDs 3HXN, 4HHB). Structural alignments of oxygenated vs. deoxygenated globins were performed in Pymol minimizing the RMSD of Cα atoms.

### Recombinant protein expression

Coding sequences for reconstructed ancestral proteins were optimized for expression in *Escherichia coli* using IDT Codon Optimization and then synthesized de novo as gBlocks (IDT). Coding sequences were cloned by Gibson assembly into vector pLIC under control of a T7 polymerase promoter (*69*). For co- expression of Ancα+Ancβ, a polycistronic operon was constructed under control of a T7 promoter and separated by a spacer containing a stop codon and ribosome binding site, as described in (*70*).

JM109 (DE3) *E. coli* cells (New England Biolabs) were heat-shock transformed and plated onto Luria broth (LB) agar plates containing 50 ug/mL carbenicillin. For the starter culture, a single colony was inoculated into 50 mL of LB with 1:1000 dilution of working- stock carbenicillin and grown overnight. 5 mL of the starter culture were inoculated into a larger 500-mL terrific broth (TB) mixture containing the appropriate antibiotic concentration. Cells were grown at 37° C and shaken at 225 rpm in an incubator until they reached an optical density at 600 nm of 0.6-0.8.

For expression of single globin proteins, 100 uM of isopropyl-β-D-1- thiogalactopyranoside (IPTG) and 25 mg/500 mL of hemin were added to each culture. Expression of single proteins in culture were done overnight at 22° C. Cells were collected by centrifugation at 4,000g and stored at −80° C until protein purification. Coexpressed proteins were induced using 500 mM IPTG expression with 25 mg/500 mL hemin for 4 hours at 37°C. Cells were collected by centrifugation at 4,000g, immediately followed by purification.

Human hemoglobin was bought commercially (Sigma-Aldridge) and resuspended in PBS.

### Protein purification by ion exchange

Purification of singly expressed ancestral proteins was performed as previously described (*26*). All singly expressed proteins (all ancestral globins except Ancα+Ancβ) were purified using ion exchange chromatography, with variation in buffer pH. All buffers were vacuum filtered through a 0.2 μM PFTE membrane (Omnipore). After expression, cells were resuspended in 30 mL of 50 mM Tris-Base (pH 6.88). The resuspended cells were placed in a 10 mL falcon tube and lysed using a FB505 sonicator (1s on/off for three cycles, each 1 minute). The lysate was saturated with CO, transferred to a 30 mL round bottom tube, and centrifuged at 20,000g for 60 minutes to separate supernatant from non-soluble cell debris. The supernatant was collected and syringe-filtered using HPX Millex Durapore filters (Millipore) to further remove debris. A HiTrap SP cation exchange (GE) column was attached to an FPLC system (Biorad) and equilibrated in 50 mM Tris-Base (pH 6.88). The lysate was passed over the column. 50 mL of 50 mM Trise-Base (pH 6.88) was run through the SP column to remove weakly bound non-target soluble products. Elution of bound ancestral Hbs was performed with 100-mL gradient of 50mM Tris-Base 1M NaCl (pH 6.88) buffer which was run through the column from 0% to 100%. For the construct Anc1-4CC, all buffer pH for purification were 6.47. 2 mL fractions were captured during the gradient process, all fractions containing red eluant were put into an Amicon ultra-15 tube and concentrated by centrifugation at 4,000g to a final volume of 1 mL. For additional purification, concentrated sample was injected into a HiPrep 16/60 Sephacryl S-100 HR size exclusion chromatography (SEC) column. The column was equilibrated in phosphate buffered saline (PBS) at pH 7.4. Purified ancestral globins elute at different volumes depending on the protein’s complex stoichiometry: 48-52 for tetramers, 56-60 for dimers, and 65-67 for monomers. The purified proteins were concentrated as mentioned above and then flash frozen with liquid nitrogen.

Coexpressed proteins Ancα + Ancβ were purified using zinc-affinity chromatography, which was performed using a HisTrap metal affinity column (GE) on a Biorad NGC Quest. Nickle ions were stripped from the column (buffer 100 mM EDTA, 100 mM NaCl, 20 mM TRIS, pH 8.0), followed by five column volumes of water. To attach zinc to the column, 0.1 M ZnSO4 was passed over until conductance was stable, approximately 5 column volumes, followed by five column volumes of water. After expression, cells were resuspended in a 50 mL lysis buffer (20 mM Tris, 150 mM Nacl, 10% glycerol (v/v), 1mM BME, 0.05% Tween-20, and 1 Roche Protease EDTA-free inhibitor tablet, pH 7.40), sonicated as described above, and the lysate passed through the prepared column. To remove non-specifically bound protein, the column was washed with 50 mL of lysis buffer. Bound protein was then eluted across a gradient of imidazole concentrations (0 to 500 mM) in a total of 100 mL elution buffer (20 mM Tris, 150 mM NaCl, 500 mM imidazole, 10% glycerol, and 1 mM BME, pH 7.4). 1 mL fractions were collected. The fraction corresponding to the second peak of UV absorbance at 280 nm has a visible red color and was collected and concentrated as described above. The concentrated solution was injected into a Biorad ENrich 650 10 x 300 columns for additional purification and eluted in PBS buffer.

### Measuring oxygen affinity and allostery

Purified proteins were deoxygenated using 10 mg/mL sodium dithionite and immediately desalted via a PD-10 column (GE Healthcare) equilibrated with 25 mL of 10 mM HEPES, 0.5 mM EDTA (pH 7.4). Eluted proteins were concentrated using Amicon Ultra-4 centrifugal filters (Millipore).

Equilibrium oxygen-binding assays were performed at 25°C using a Blood Oxygen Binding System (Loligo Systems) with 0.1 mM protein (heme concentration) dialyzed in 100 mM HEPES, 0.5 mM EDTA buffer. Protein solutions were sequentially equilibrated at 3–5 oxygen tensions (PO₂), achieving 30–70% saturation, while continuously monitoring absorbance at 430 nm (deoxy peak) and 421 nm (oxy/deoxy isosbestic point). Fractional saturation was plotted against PO₂, and the Hill equation was fitted to each dataset to estimate P₅₀ (PO₂ at half- saturation) and the Hill coefficient (n₅₀, slope at half-saturation). Confidence intervals (95%) were calculated by multiplying the standard error of the mean from replicate experiments by 1.96.

To assess potential allosteric regulation by organic phosphate effectors, assays were conducted under three conditions: without effectors (stripped), with 0.5 mM IHP, and with 0.5 mM ATP. Most experiments were performed with IHP due to its stronger allosteric effect compared to ATP and their qualitatively similar modulation of Hb-O₂ affinity (*22, 71*). Although IHP may not have been the physiological effector in ancestral organisms, it has been shown to allosterically regulate hemoglobins across major vertebrate lineages, whereas effectors such as 2,3- bisphosphoglycerate (BPG), ATP, and GTP exhibit lineage-specific effects. Thus, IHP serves as a general polyanion for assessing the allosteric capacity of ancestral Hb, a well-established approach in hemoglobin studies (*25, 72*).

### Fluorescence emission scan

Purified proteins were deoxygenated with 10 µM sodium dithionite to remove oxidized hemoglobin, then desalted using a PD-10 column (Cytiva) equilibrated with PBS. If necessary, eluted proteins were concentrated using Amicon Ultra-15 filters prewashed with PBS.

To evaluate conformational states as a function of oxygenation, proteins were aliquoted into two 150 µL samples at 150 µM concentration. Buffer-only aliquots were prepared for fluorescence background correction. Oxygenated hemoglobin samples were verified by UV-Vis spectroscopy, confirming the characteristic double peak at 480 nm. Samples were transferred into a sealed sub- micro quart fuorometer cell (Starna Cells, Inc.). Fluorescence scans were performed using a Fluorolog-3 (Horiba) with excitation at 280 nm and emission recorded from 307–500 nm in 1 nm increments. Slit widths for excitation and emission ranged from 4–7 nm. To mitigate inner-filter effects of hemoglobin, a front-facing mirror setup was used, an established approach for Hb fluorescence studies (*73*).

For deoxygenated samples, each aliquot was incubated in a custom anaerobic chamber saturated with nitrogen for two hours, with periodic resuspension to facilitate oxygen exchange. At the time of measurement, the sample was transferred to the same sealed sub-micro quartz fluorometer cell used in oxygenation condition. Fluorescence scans were conducted immediately after chamber removal.

### Molecular dynamics simulations

Models of Anc1 and Anc1-4CC were generated using the AlphaFold3 web server (*74*). Multimeric predictions were carried out to model the quaternary structure by including four copies of each monomer, and heme cofactors (deoxygenated) were included using the ligand modeling capabilities of AlphaFold3. Model confidence was assessed via the predicted template modeling (pTM) and interface predicted template modeling (ipTM) scores. Anc1 and Anc2 exhibited pTM scores of 0.96 and 0.92, respectively, and ipTM scores of 0.95 ad 0.91, respectively, indicating high confidence in predictions of tertiary and quaternary structures.

The VMD psfgen, solvate, and autoionize plugins were used to build all systems, and all analysis was done using VMD with in-house Tcl scripts or MDAnalysis (*75, 76*). Water boxes were made large enough to ensure no interactions occurred between periodic images. For both systems, Cl^-^ atoms were added to neutralize charge. The N-terminal residue of all proteins was capped with – NH^3+^ and the *C*-termini with a –COO^−^ carboxylate group. Simulations were run using NAMD 2.144 with the CHARMM36^5^ force field and the TIP3P explicit model of water (*77, 78*). A cutoff of 12 Å was used for van der Waals interactions, and long-range electrostatic forces were computed using the Particle Mesh Ewald method with a grid point density of >1 Å−3. The SHAKE algorithm was used with a timestep of 2 fs. The NpT ensemble was used at 1 atm with a hybrid Nosé-Hoover Langevin piston method, a 200-fs decay period, and with a 100-fs damping time constant. Temperature was set to 300 K. All systems were minimized for 2000 steps, followed by 100 ps of simulation with hemes and backbone atoms constrained, after which constraints were lifted for production MD.

Previous experiments have shown that Cl^-^ can act as an allosteric effector of hemoglobin (*79, 80*). Because of this, and because the Cl^-^ is necessary to neutralize the excess negative charge of the overall system, the collective variables module in NAMD was used in simulations without IHP to prevent Cl^-^ binding to interfacial regions. A harmonic potential of 100 kcal/mol was applied to any Cl^-^ that drifts within 15 Å of any residue at the α1α2 and β1β2 interfaces. Beyond this cutoff, no potential was applied. This approach permits a charge neutral system, without affecting the allosteric dynamics of the protein in an unintended fashion. Initial equilibration simulations of Anc1 and Anc1-4CC were initiated from this setup, with no further modifications.

Systems with IHP were prepared by first aligning the H-helices of the Anc1 or Anc1-4CC chains 2 and 3 to the H-helices of the beta chains of the human hemoglobin bound to IHP (PDB code 3HXN). The IHP molecule was then saved and combined with the Anc1 or Anc2 structure, with subsequent solvation and ionization. Parameters for IHP were generated using the CGenFF web server, which resulted in a param penalty of 6.5 and a charge penalty of 7.5, indicating high quality predictions (*81*).

To initiate simulations with and without oxygen after removing IHP, the coordinates of the protein and hemes, but without IHP, were saved from the final frame of the Anc1-4CC with IHP simulation. This was solvated, ionized, and minimized as above. VMD’s mutate plugin was used on the saved structure, prior to solvation, to revert the four CC residues to their amino acid state in Anc1, which was then solvated, ionized, and minimized as above. These two systems were used to initiate production MD for Anc1-deoxy and Anc1-4CC-deoxy starting from a putative T-like state. To prepare identical systems in the oxy state, the same saved structure from the Anc1- 4CC with IHP simulation as above was used but with the PLO2 patch added to the heme molecules during psf generation, followed by solvation and ionization as above. This resulted in a total of four IHP-removed systems: Anc1-4CC-oxy, Anc1-4CC-deoxy, Anc1-oxy, and Anc1- deoxy. Simulations were run identically as in the equilibration simulations as described above.

To prepare the Anc1-t142S system, an initial structure was predicted, solvated, and ionized as described above, using the PLO2 patch during psf generation to add O2 ligands to the heme molecules. This structure was used to run a 100 ns equilibrium simulation using identical parameters as above. Hemoglobin in the final frame of this simulation was saved and used to dock IHP into. Hydrogens were added, Gasteiger charges were calculated, and atom type and torsion groups were assigned for IHP using autodocktools. AutoDock Vina^9^ was then run with the entire protein used as a search space. Four of the top five lowest energy docked structures were found in the central cavity (*82*). The second lowest energy structure (−6.3 kcal/mol) was selected for further simulation and saved as a structure together with the hemoglobin as the starting point for simulation (Fig. 5A). As a control, VMD’s mutate plugin was used on this structure to mutate the sequence of hemoglobin to Anc1. Both structures were solvated and ionized as above, and simulations were run identically as in the equilibration simulations as described above. All distributions of conformational metrics were calculated by excluding the first 20 ns after equilibration.

## Acknowledgements

We thank members of the Thornton and Storz labs for helpful advice and comments. Supported by NIH R35GM145336 (JWT), NIH R01GM139007 (JWT), NIH R01HL159061 (JFS), and NSF IOS-2114465 (JFS).

