## Supplementary Figures S1-28 for "Origins of allostery in vertebrate hemoglobin evolution"

Carlos R. Cortez-Romero *et al.*

This PDF includes:

Figs. S1 – S28

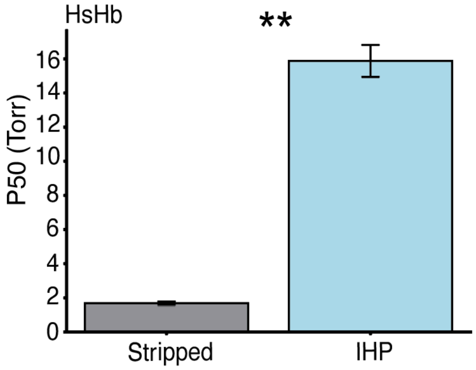


**Fig S1. Oxygen affinity of human hemoglobin in the presence and absence of allosteric effector.** IHP was added at 500 μM. Error bars show standard error of measurement for 3 replicates. **, $P$<0.01 between conditions via Welch's two-sample t-test.

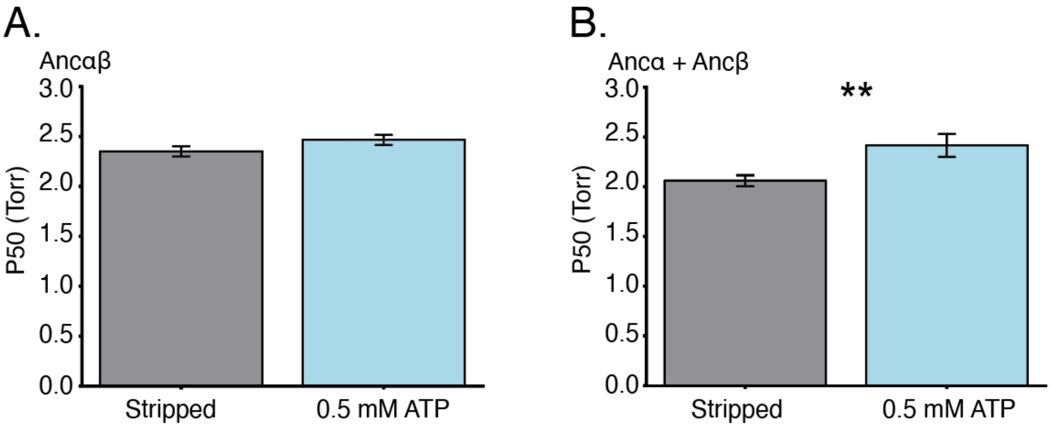


**Fig S2. Allosteric response of ancestral proteins to ATP. (A)** Oxygen affinity of Ancαβ in the presence and absence of ATP. In grey, stripped medium; blue, 500 μM of ATP is added. Error bars, standard error of measurement. **(B)** Oxygen affinity of Ancα + Ancβ, n = 10. **, $P$<0.01 between conditions via Welch's two-sample t-test.

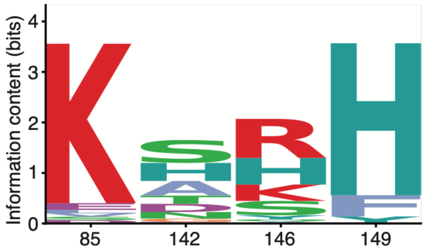


**Fig S3. Conservation of central cavity positions in extant hemoglobins.** Logo plot of per-position information content (bits) calculated from a multiple sequence alignment of vertebrate hemoglobin β-subunit sequences at the four central cavity residue positions that were substituted between Anc1 to Anc1-4CC. Maximum information content is 4.32 bits (invariant position).

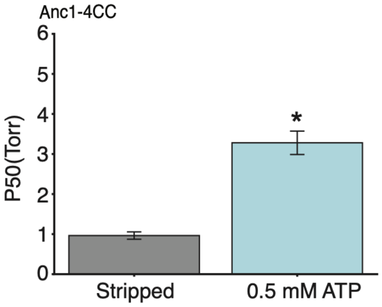


**Fig S4. Allosteric response to ATP in Anc1-4CC.** Bar graph of oxygen affinity of Ancαβ in the presence and absence of ATP. In grey, stripped condition, where no ATP is in solution. In blue, ATP condition, where 500 µM of ATP is added to solution. Error bars represent standard error of measurement, n = 3. *, $P<0.05$ between conditions via Welch's two-sample t-test.

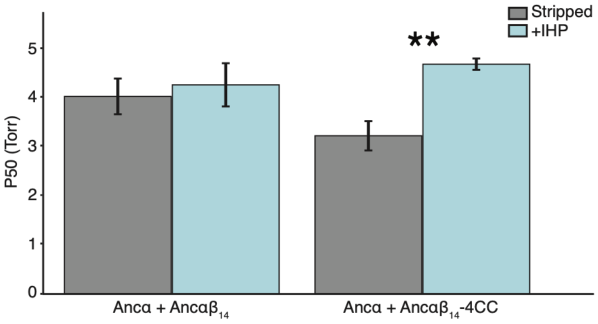


**Fig S5. Central cavity mutations are robust to heteromerization.** A bar graph of Ancα + Ancαβ$_{14}$, a known stable heterotetramer, without and with the central cavity mutations (Ancα + Ancαβ$_{14}$ -4CC) in the presence and absence of IHP. In grey, stripped condition, where no IHP is in solution. In blue, IHP condition, where 500 µM of IHP is added to solution. Error bars represent standard error of measurement, n = 3. ** represent FDR  <0.05 between conditions via Welch's two-sample t-test and Benjamini-Hochberg FDR procedure.

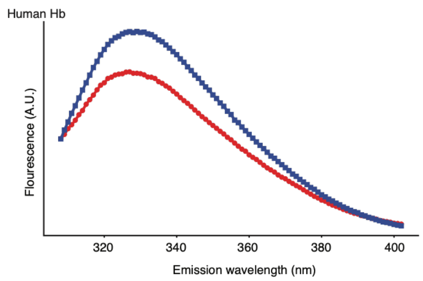


**Fig S6. Conformational heterogeneity of human Hb.** Fluorescence emissions scans of Human Hb when excited at 280 nm. In red, protein is oxygenated; in blue, deoxygenated. Error bars represent standard error of measurement, n = 10.

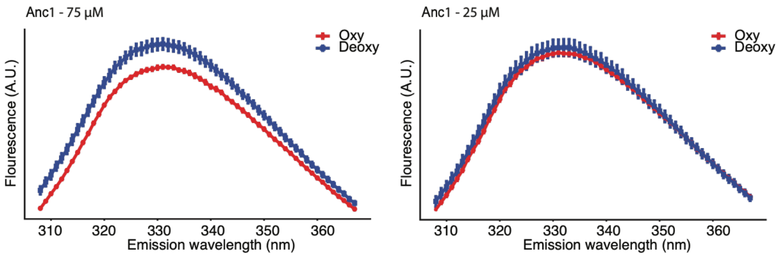


**Fig S7. Concentration-dependent fluorescence signal of conformational change in Anc1 indicates tetramer-dependence.** Fluorescence emissions scans of Anc1-4CC when excited at 280 nm, at two protein concentrations. In red, protein is oxygenated; blue, deoxygenated. Error bars represent standard error of measurement, n = 10. As expected, the difference between conditions increases with concentration, because occupancy of the tetrameric stoichiometry increases with concentration, and quaternary heterogeneity requires tetramerization. The concentrations used are in the range at which occupancy increases with concentration (see ref. 26).

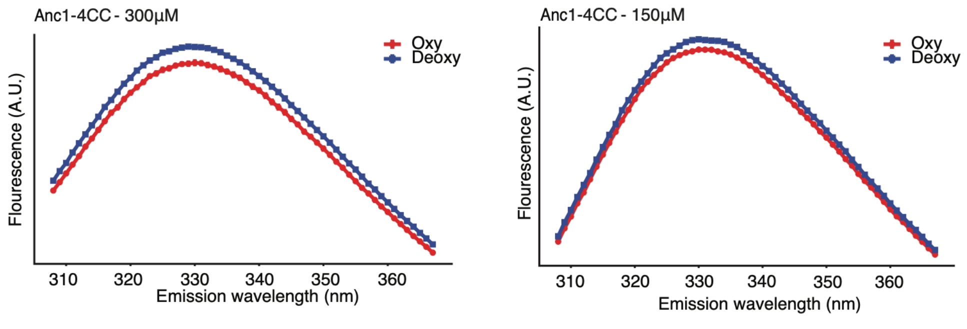


**Fig S8. Concentration-dependent fluorescence signal of conformational change in Anc1-4CC indicates tetramer-dependence.** Fluorescence emissions scans of Anc1-4CC when excited at 280 nm, at two protein concentrations. In red, protein is oxygenated; blue, deoxygenated. Error bars represent standard error of measurement, n = 10. As expected, the difference between conditions increases with concentration, because occupancy of the tetrameric stoichiometry increases with concentration in this range (see fig. S9), and quaternary heterogeneity requires tetramerization.

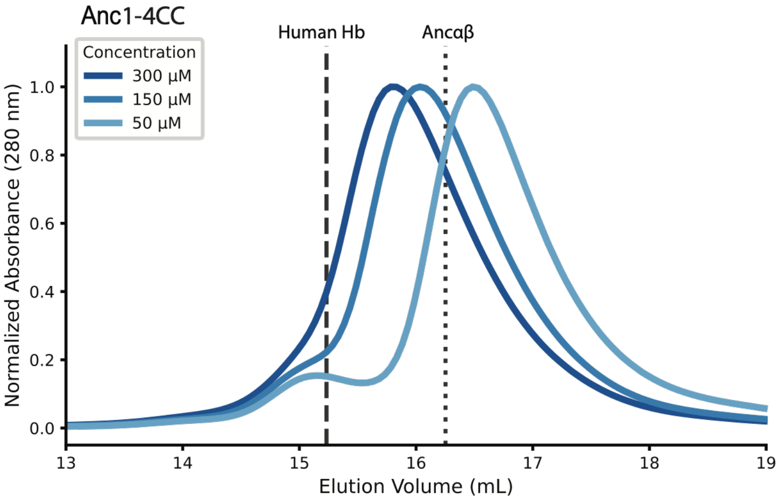


**Fig S9. Size exclusion chromatography elution of Anc1-4CC indicates increased tetramer assembly at higher concentrations.** Elution traces of size exclusion chromatography at three protein concentrations. Lines at the elution peaks of Human Hb, a known tetramer, and Ancαβ, a known, dimer shown are shown as reference.

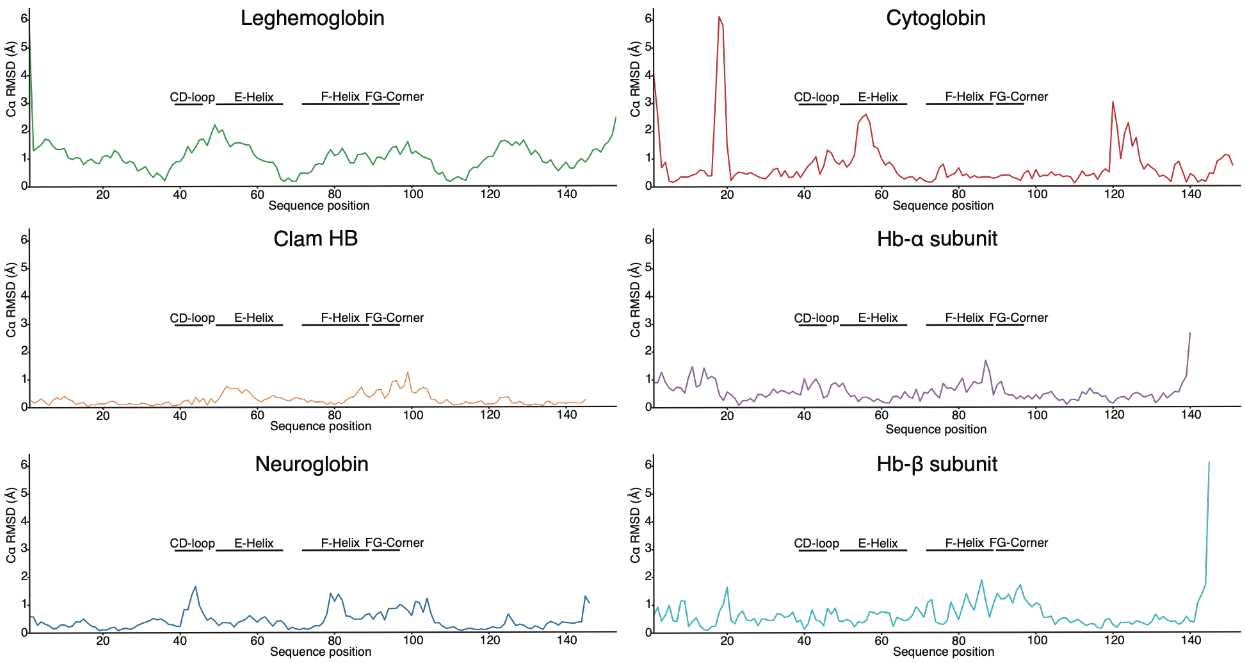


**Fig S10. Individual per-residue Cα deviation between oxygenated and deoxygenated crystal structures of globin proteins.** Each color represents a distinct globin protein family member: leghemoglobin (green), clam hemoglobin (red), neuroglobin (dark blue), cytoglobin (teal), Hb α subunit (purple), Hb β subunit (cyan). For each, the crystal structures of oxy and deoxy structures of the same protein were aligned to minimize RMSD across Cα atoms, and the distance between Cα atoms of the two structures at each residue are plotted across the length of the protein. PDB IDs are in the legend for main text Fig. 4.

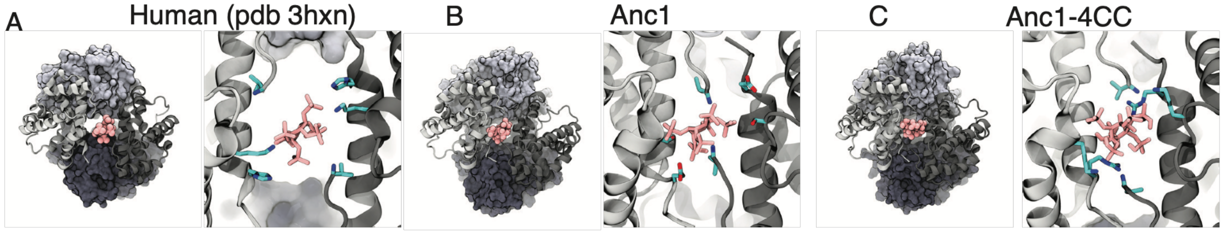


**Fig S11. IHP binding cavity of human Hb and modeled ancestral hemoglobin precursors.** **(A)** Surface representation (left) and close-up view of the central cavity (right) of Human Hb (PDB: 3HXN) with IHP (pink). Key cavity-lining residues are shown as cyan sticks. Surface representation, alpha subunits; cartoon, beta subunits. **(B)** Equivalent views for Anc1. **(C)** Equivalent views for Anc1-4CC. Structural models for panels B and C were generated by AlphaFold3 and docked with IHP as described in Materials and Methods. Structures shown in B and C are before molecular dynamics simulations.

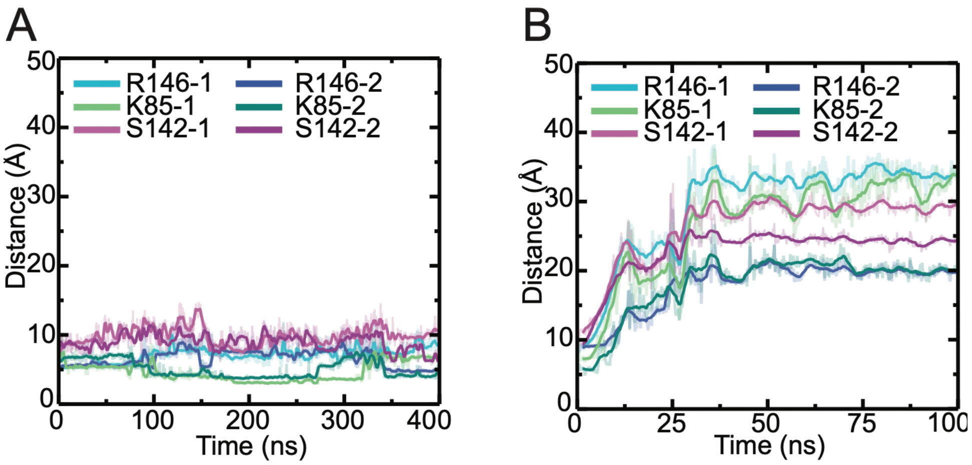


**Fig S12. Time course of molecular dynamics simulation measuring the distance of central cavity residues to IHP phosphate in Anc1-4CC and Anc1.** Distances between the center of mass of IHP and central cavity residues in Anc1-4CC **(A)** or Anc1 **(B)** across each MD simulation. Indices 1 or 2 distinguishes the two copies of each residue in the symmetrical cavity. The distances in panel B become large as IHP flies out of the pocket (see fig. S13).

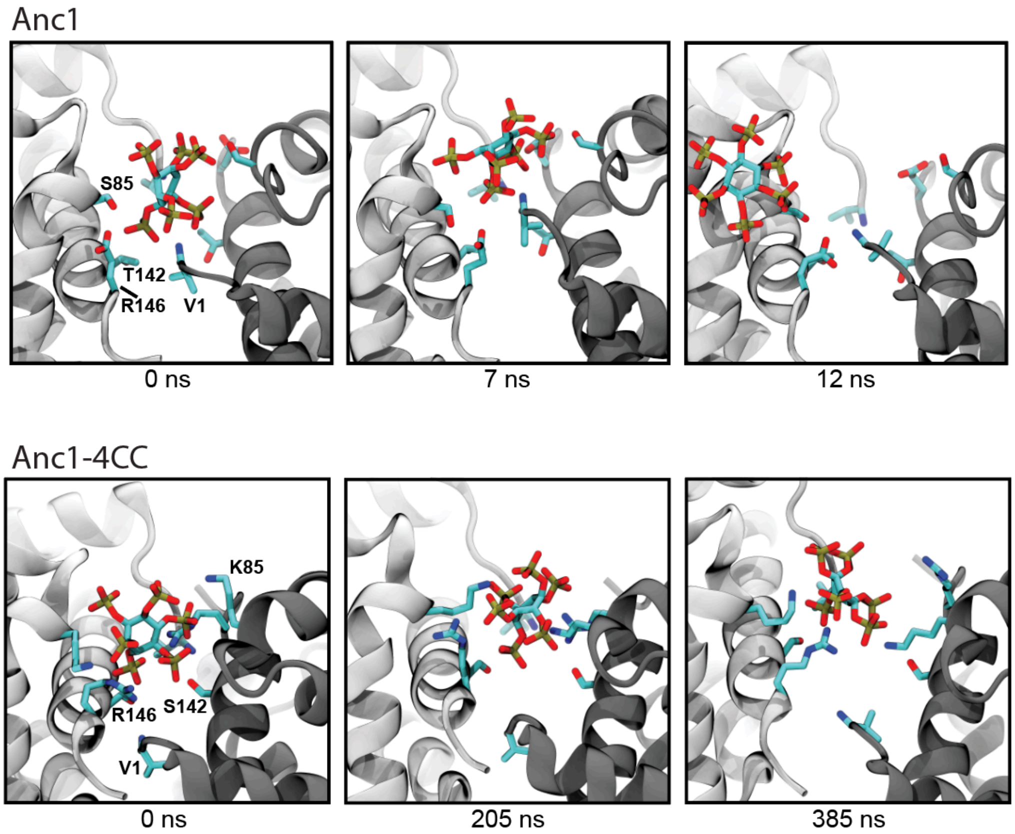


**Fig S13. Structural view of IHP binding in Anc1 and Anc1-4CC.** Snapshots from MD simulations showing the central cavity residues interacting with IHP (shown as pink sticks) at the indicated time points. *(Top)* Anc1 at 0, 7, and 12 ns showing residues s85, t142, r146, and the Val1 N-terminus in cyan. *(Bottom)* Anc1-4CC at 0, 205, and 385 ns showing the substituted residues (K85, S142, and R146) and Val1. IHP rapidly dissociates from the Anc1 central cavity, while Anc1-4CC maintains persistent contacts throughout the simulation.

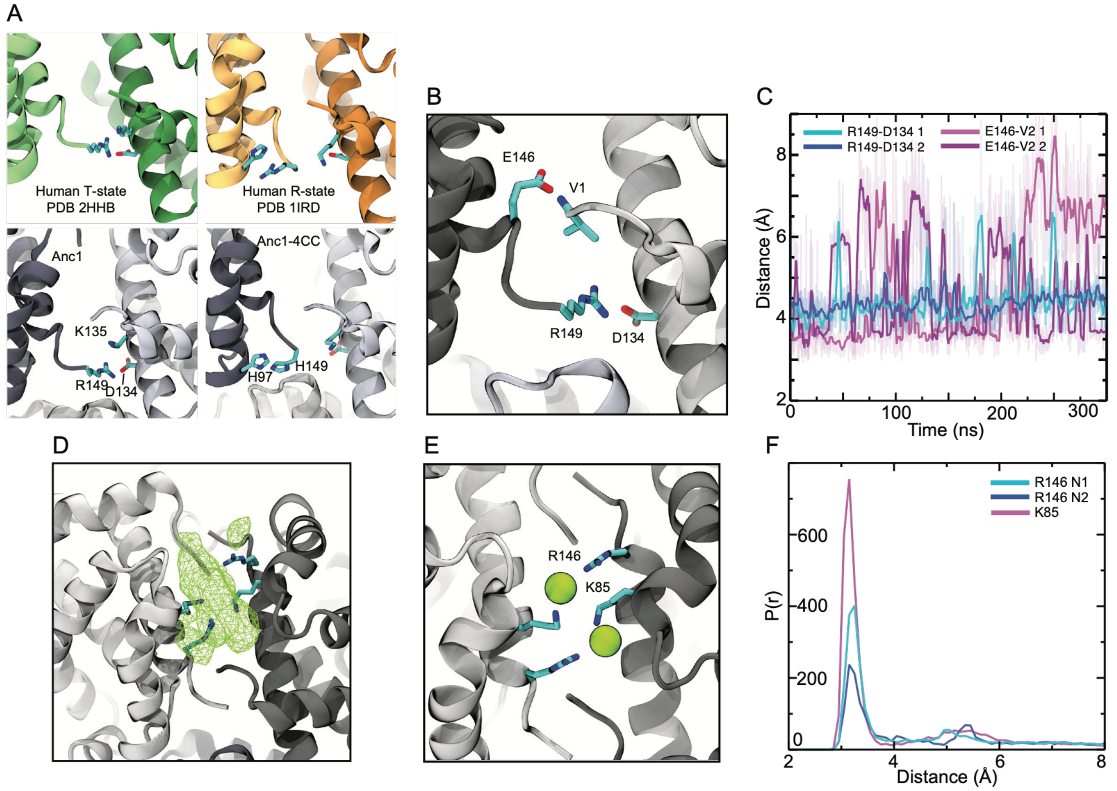


**Fig S14. Effects of substitutions on the electrostatic environment of the central cavity in the absence of IHP. (A)** Structural comparison of the central cavity region of human Hb in the T and R states, as well as Anc1 and Anc1-4CC, all without IHP. Interaction of substituted residue r149H with D134 within the central cavity (in Anc1 and the human T state) or with H97 (in Anc1-4CC and the R-state) is shown. **(B)** Close view of the Anc1 central cavity without IHP showing interactions of e146 with Val1 (V1) and Arg149 with Asp 134 (D134); **(C)**, distance traces for R149–D134 (cyan) and E146–V2 (pink) contact pairs across the central cavity. Indices 1 or 2 refer to the two iterations of each residue in the symmetrical cavity. **(D)** Central cavity of Anc1-4CC in the absence of IHP. Surface representation showing the electrostatic environment and unoccupied space in the central cavity (green mesh). Sticks, R146 and K85. **(E)** R146 and K85 coordinate chloride ions (green spheres) throughout the trajectory. **(F)** Probability distribution across the simulation of the distances from atoms on each of these residues to the nearest chloride atom.

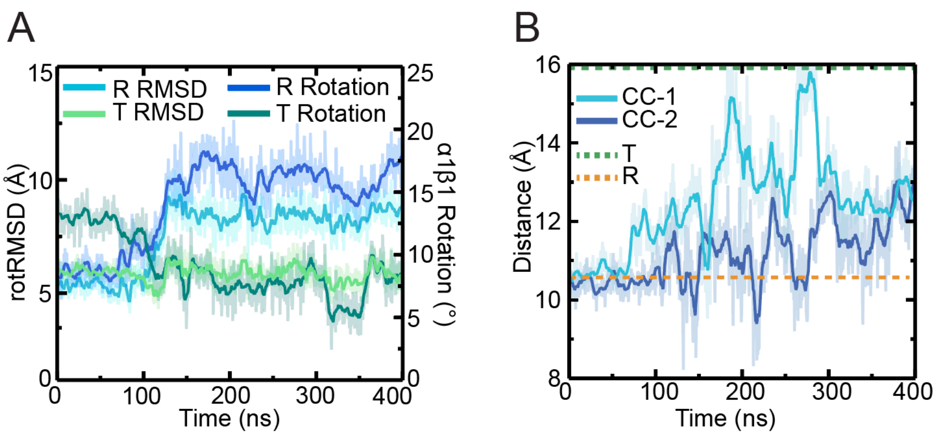


**Fig S15. Quaternary similarity across the MD simulation of Anc1-4CC with IHP to human Hb T and R states. (A)** Angle of rotation and the alpha-carbon RMSD relative to the human T and R conformations (2HHB and 1IRD) is shown for the simulated complex at each time point. **(B)** Distance measurement between the H-helices of two subunits across the central cavity at each timepoint. Indices refer to the two central cavities in the homotetramer. For all metrics, a transition is evident during the first 100 ns from a more R-like starting point (which was generated without IHP) to a more T-like conformation.

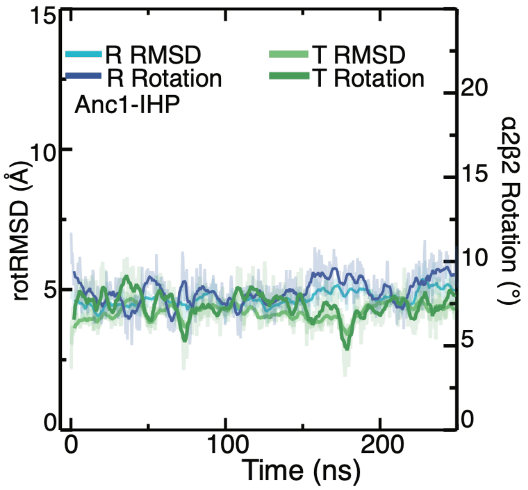


**Fig S16. Anc1+IHP does not undergo an R-to-T-like quaternary transition.** Rotational RMSD (rotRMSD, Å, left axis) and dimer rotation angle (right vertical axis) of Anc1+IHP relative to human Hb in the oxygenated R or deoxygenated T conformation. The rotation angle represents the angle between the central axes of the unaligned dimers ($\alpha_2\beta_2$) when two tetramers are aligned using the alpha-carbons in the other dimer ($\alpha_1\beta_1$). Compare to fig. S15, which shows the R-to-T transition of Anc1-4CC in the presence of IHP.

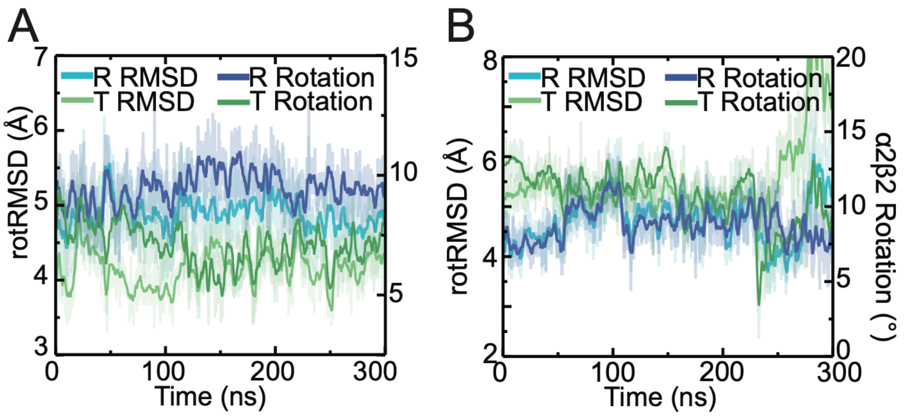


**Fig S17. Comparison of quaternary structure of Anc1 and Anc1-4CC in the absence of oxygen and IHP. (A)** Rotation angle and rotRMSD of Anc1 relative to human R-state (blues) and T-state (greens). **(B)** rotation angle and rotRMSD of Anc1-4CC relative to human R-state (blues) and T-state (greens). Anc1 favors the T conformation, whereas Anc1-4CC does not.

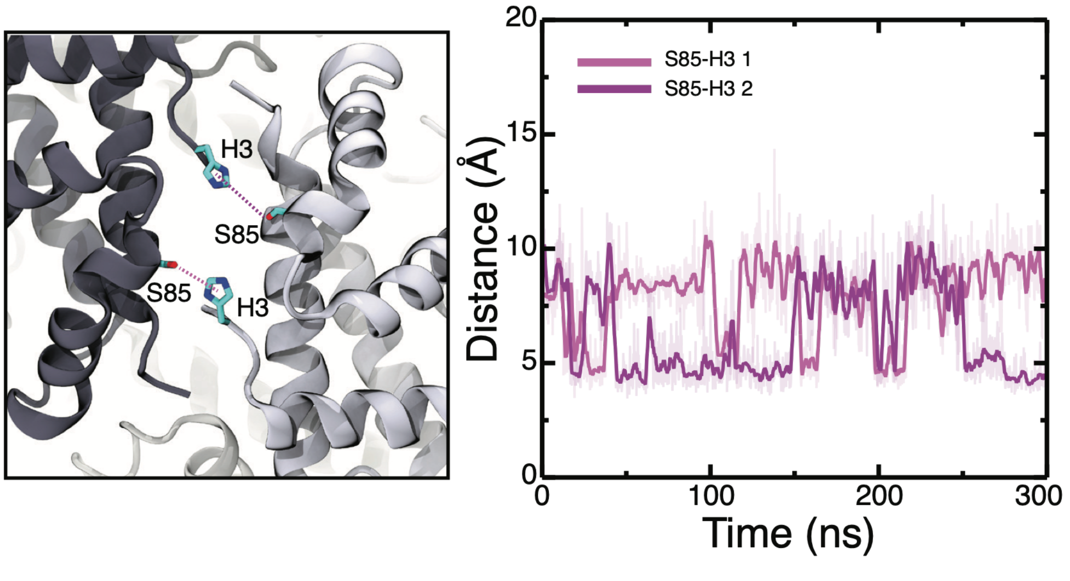


**Fig S18. Ser85 makes contacts with His3 across the central cavity in Anc1.** *(Left)* Representative structural snapshot from MD simulation of Anc1 without IHP showing ancestral residue ser85 (cyan) on one subunit forming contacts (dashed pink lines) with residues in H3 of the opposing subunit. *(Right)* Distance (Å) between S85 and His3 contact atoms, measured in both isologous iterations across the central cavity.

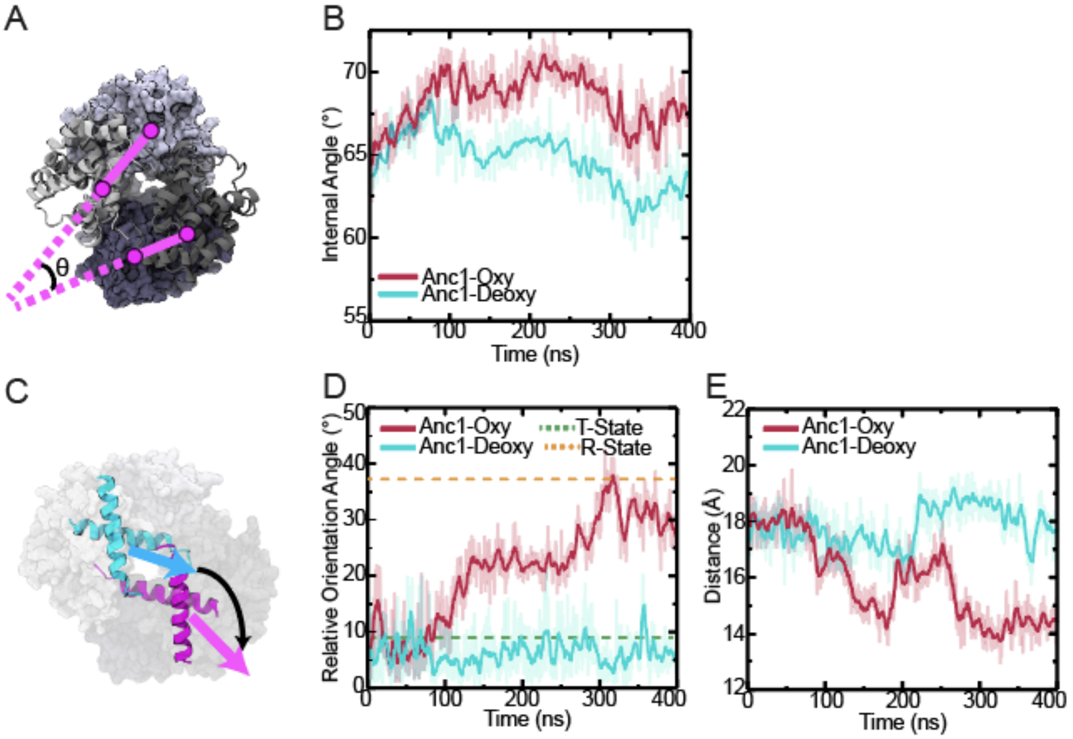


**Fig S19. Oxygen-dependent conformational heterogeneity of Anc1. (A)** Schematic representation of measuring the internal rotation angle. Dots indicate the center of mass of each subunits. Lines, the axis of each dimer in the tetramer, generated by connecting the centers of the two subunits in each dimer. θ, angle between the two axes in a tetramer is the internal rotation angle. **(B)** Internal rotation angle of oxygenated Anc1 and deoxygenated Anc1 across their trajectories. **(C)** Schematic representation of measuring the relative orientation angle of two H-helices. Colored arrows indicate the chosen principal axis of the H-helices within a dimer, and the black arrow indicates the rotation required to align the axes from the two dimers to each other. Dotted lines, this metric for crystal structures of human Hb in oxy and deoxy conformations (1IRD and 2HHB). **(D)** Relative orientation angle across the trajectory of oxygenated Anc1 and deoxygenated Anc1. **(E)** Proximity of the CD loop to its neighboring subunit across IF2, measured as the average distance between the Cα of H47 on the CD loop of one subunit and K98 on the FG corner of the other.

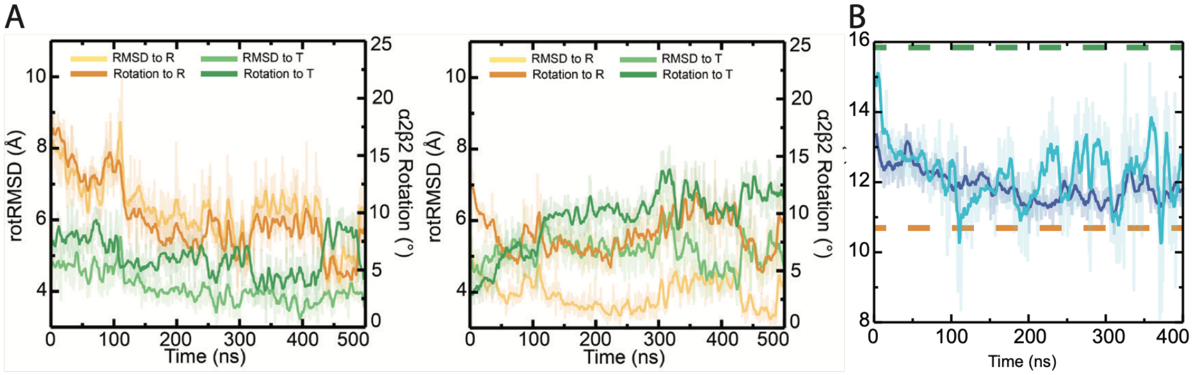


**Fig S20. Oxygen dependent conformational change in Anc1. (A)** (*Left)* For oxygenated Anc1, the rotational RMSD (rotRMSD, Å, *left vertical axis*) and dimer rotation angle (*right vertical axis*) are shown when aligned to human Hb in the oxygenated R or deoxygenated T conformation. The rotation angle represents the angle between the central axes of the unaligned dimers (corresponding to $\alpha_2\beta_2$ in human Hb) when two tetramers are aligned using the alpha-carbons in the other dimer ($\alpha_1\beta_1$). *(Right)* The same metric is plotted for $\alpha_2\beta_2$ when the $\alpha_1\beta_1$ dimers are aligned. **(B)** Distance between the H helices across the central cavity (Å) for oxygenated Anc1 over time; dashed orange line indicates the T-state reference distance (2HHB). Dark and light blue series represent the two pairs of H helices in the homotetramer. For all metrics, a transition to a more R-like state is apparent.

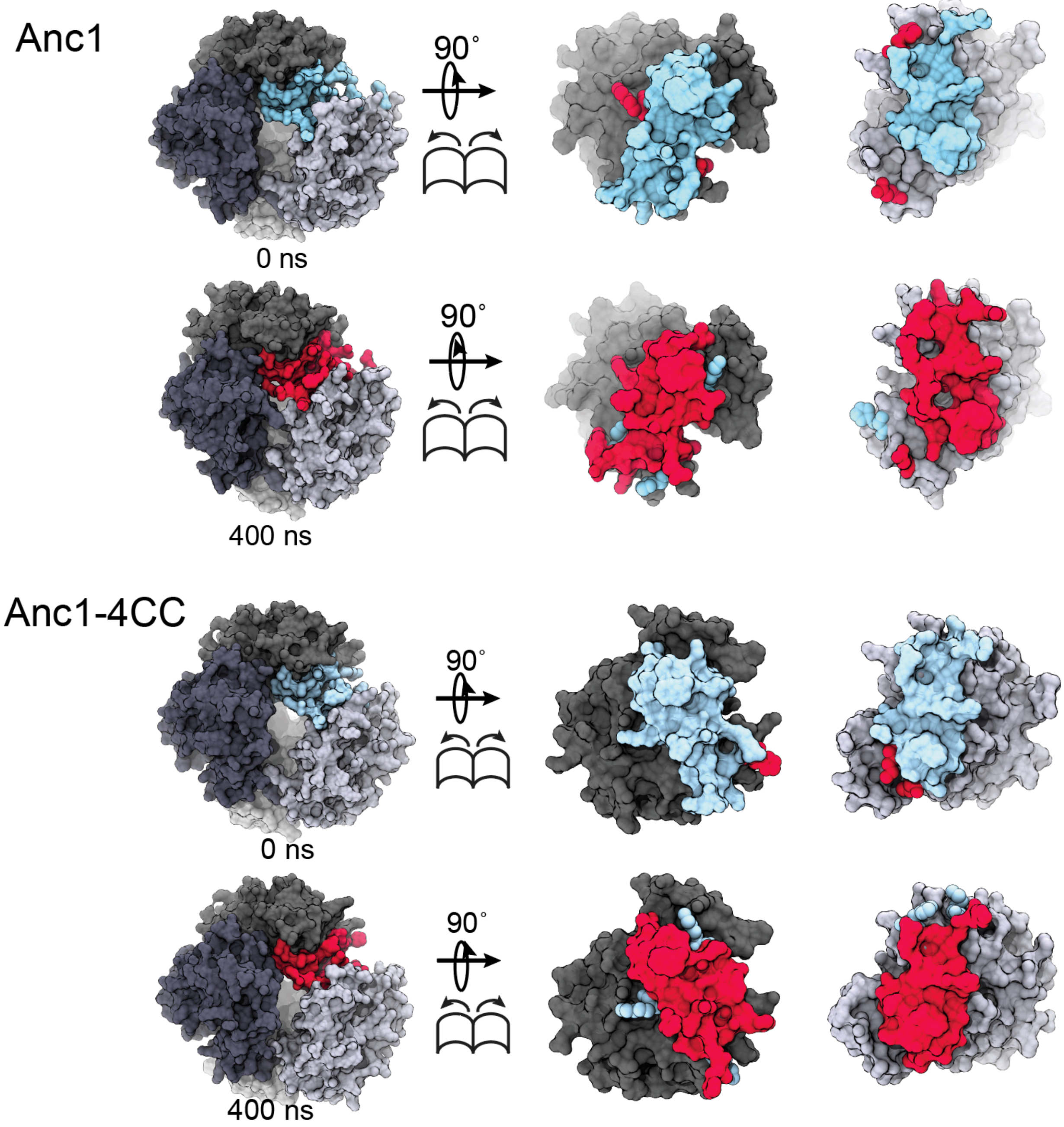


**Fig S21. Changes in the tetramerization surface of Anc1 and Anc1-4CC caused by oxygen-binding.** Surface representations of the tetramerization interface in Anc1 *(top two rows)* and Anc1-4CC *(bottom two rows)*. Surface residues buried in the interface are colored blue at 0 ns and red at 400 ns in the trajectory with oxygenated heme. The oxygenated trajectories were initiated using the endpoint structures of trajectories with deoxygenated heme, so changes from 0 to 400 ns illustrate the transition induced by oxygenation. *(Left)* intact tetramers, *(Right)* each tetramer split into its component dimers. To facilitate comparison, the buried surface at both timepoints is shown in both representations, but with the 0 ns surface in front (first series in each group) or the 400 ns surface in front (second series).

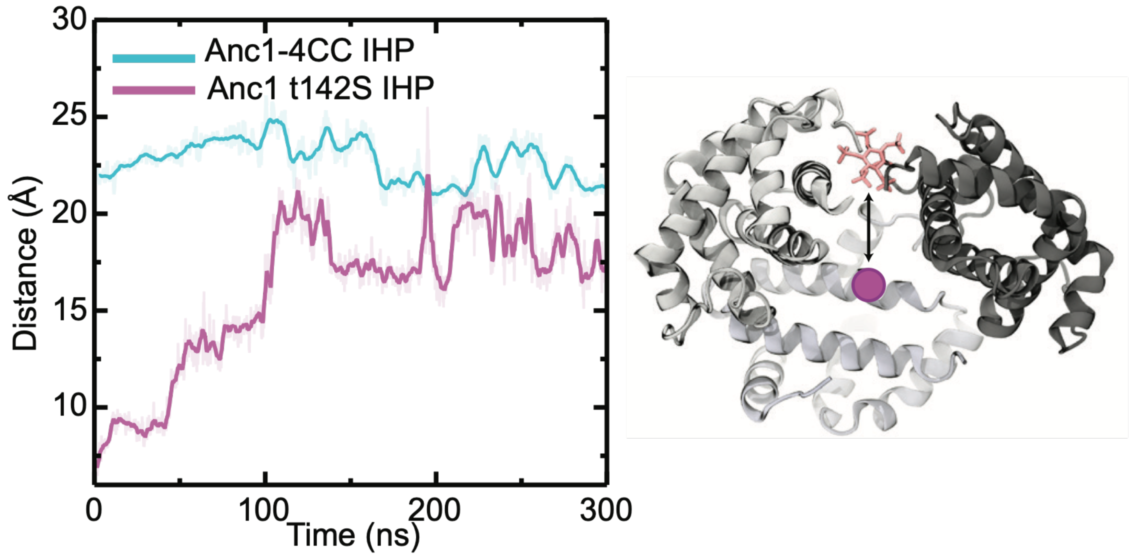


**Fig S22. IHP occupies a location deeper in the central cavity of Anc1-t142S relative to Anc1-4CC.** *(Left)* Distance between the center of mass of the heterotetramer and the center of mass of IHP across the trajectories of Anc1-4CC- IHP and Anc1-t142S. After ~100 ns, an equilibrium is reached, and interaction with S142 keeps IHP lower in the cavity than in Anc1-4CC. *(Right)* Schematic of this measurement, with one subunit removed for clarity.

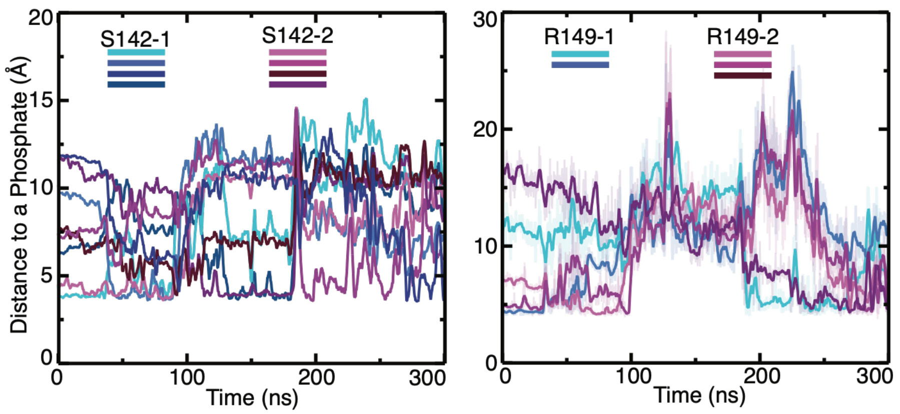


**Figure S23. MDS trajectory of distances between S142 and R149 to IHP.** Distances between phosphate atoms of IHP and S142 (Oγ atom, *Left*) or R149 (Cζ atom, *Right*) are shown throughout the simulation. The indices 1 or 2 indicate the copies of each residue in the symmetrical cavity. The multiple shades represent distance to different phosphates on IHP. At virtually all timepoints, each residue is interacting one phosphate.

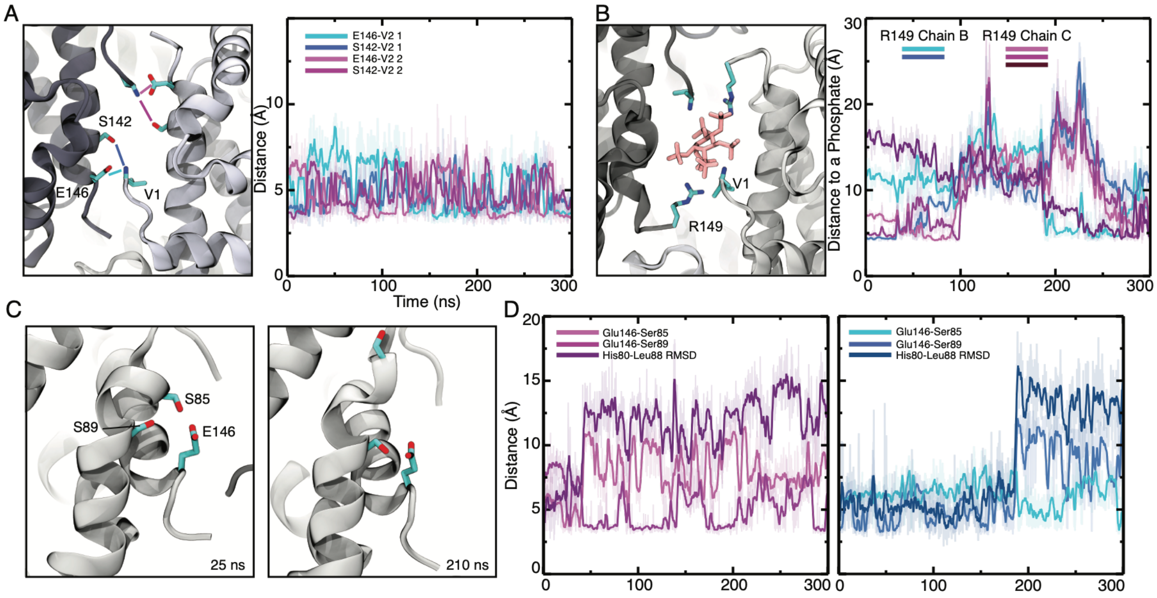


**Fig S24. IHP coordination by Anc1-t142S. (A)** Structural views of the second central cavity between subunits 1 and 4 of Anc1-t142S, where IHP is not bound. Stable contacts are formed between Val1 (V1) and S142 or E146 residues; distances between these residues are plotted across the trajectory. **(B)** Structural views of the Anc1-t142S central cavity, β domains, where IHP binds. with IHP *(Right)*. Distance from the Cζ atom of Arg149 (chains B and C) to the nearest IHP phosphate oxygen over 300 ns; colored bars at top indicate time periods of persistent phosphate contact *(Left)*. **(C)** Snapshots at 25 ns and 210 ns showing Glu146 in contact with Ser85 and Ser89, which induces conformational changes in the F-helix. **(D)** Distance traces for Glu146–Ser85 and Glu146–Ser89 interactions overlaid with His80–Leu88 RMSD over 300 ns for two independent β-subunit chains (*Left and Right panels*). Increase in RMSD is correlated with the loss of the Glu146-Ser85 interaction and gain of the Glu146-Ser89 interaction, illustrating coupling between C-terminal orientation and the structure of the F-helix.

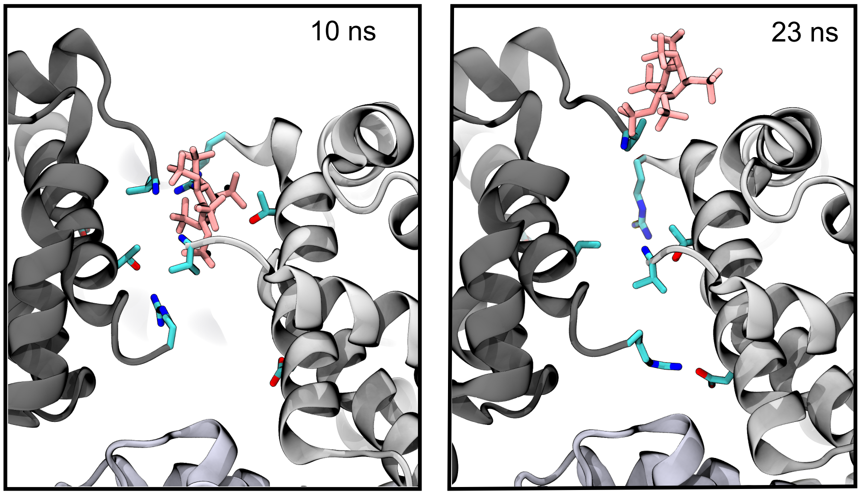


**Fig. S25. IHP is ejected from the central cavity of Anc1 pocket when it is docked in the same location as in Anc1-t142S.** Two timepoints are shown from the simulation of Anc1 with IHP, which was initiated based on the same starting structure as used for Anc1-t142S with oxygenated heme, but with residue 142 mutated to the ancestral threonine.

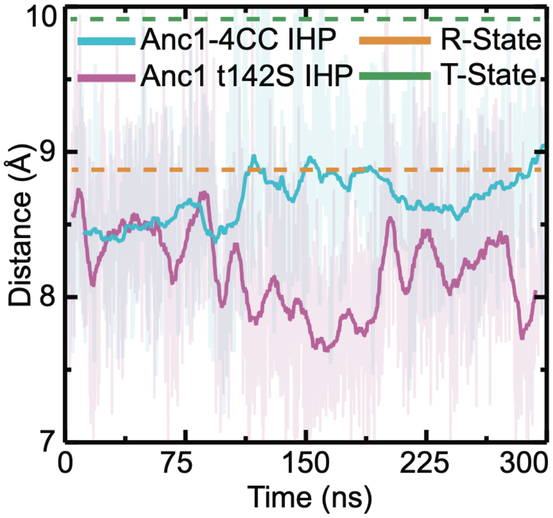


**Fig S26. Packing in the heme binding pocket of Anc1-t142S with oxygen.** The helix E-helix F distance (mean across the tetramer's four subunits) is plotted across simulations, as shown in Fig. 5D,E. Dotted lines, this distance for human R- and T-state crystal structures (PDB 1IRD and 2HHB).

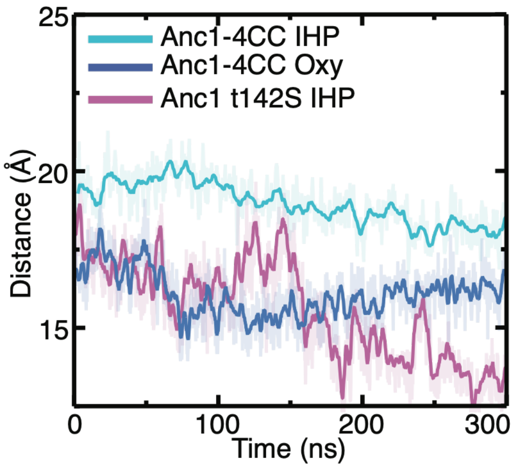


**Fig S27. IHP binding alters conformation at the tetramerization interface of Anc1-t142S with oxygen.** The distance from the CD loop to the FG corner across the interface is shown (mean across all four iterations in the homotetramer) as shown in Fig 5F,G.

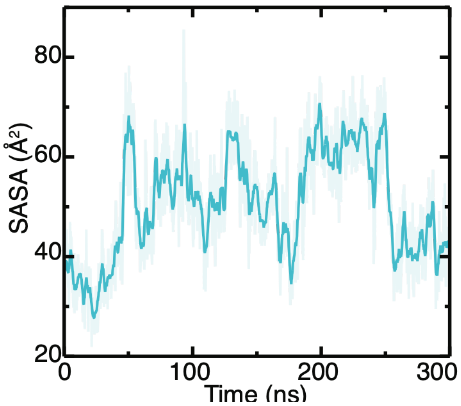


**Fig S28. IHP increases solvent-accessibility of IF2 in Anc1-t142S.** Increase in average solvent-accessible surface area of Trp40 residues across this simulation.
